# LINE-1 and the cell cycle: protein localization and functional dynamics

**DOI:** 10.1101/147587

**Authors:** Paolo Mita, Aleksandra Wudzinska, Xiaoji Sun, Joshua Andrade, Shruti Nayak, David J. Kahler, Sana Badri, John LaCava, Beatrix Ueberheide, Chi Y. Yun, David Fenyo, Jef D. Boeke

## Abstract

LINE-1/L1 retrotransposon sequences comprise 17% of the human genome. Among the many classes of mobile genetic elements, L1 is the only autonomous retrotransposon that still drives human genomic plasticity today. Through its co-evolution with the human genome, L1 has intertwined itself with host cell biology to aid its proliferation. However, a clear understanding of L1’s lifecycle and the processes involved in restricting its insertion and its intragenomic spreading remains elusive. Here we identify modes of L1 proteins’ entrance into the nucleus, a necessary step for L1 proliferation. Using functional, biochemical, and imaging approaches, we also show a clear cell cycle bias for L1 retrotransposition that peaks during the S phase. Our observations provide a basis for novel interpretations about the nature of nuclear and cytoplasmic L1 ribonucleoproteins (RNPs) and the potential role of DNA replication in L1 retrotransposition.

## INTRODUCTION

Retrotransposons are genetic elements that move within the host genome through a “copy and paste” process utilizing an RNA intermediate. 17% of the human genome is made up of copies of the Long Interspersed Nuclear Element-1 (LINE-1 or L1) retrotransposon. L1 is the only autonomous human retrotransposon still able to “jump” through a process called retrotransposition. The full length, active L1 retrotransposons consists of a 5’ untranslated region (UTR) containing a bidirectional promoter, two open reading frames (ORFs), ORF1 and ORF2, separated by a short inter-ORF region and a 3’ UTR with a weak polyadenylation signal (Doucet et al., 2015). Upon transcription by RNA polymerase II, the 6 kb bicistronic L1 mRNA is poly-adenylated and exported into the cytoplasm. LINE-1 exhibits “cis-preference” (Wei et al., 2001), the mechanism of which is unknown. A popular model posits that newly translated ORF1 and/or ORF2 proteins preferentially bind the mRNA molecule that encoded them (Boeke, 1997). Interestingly, translation of the second ORF, ORF2p, is mediated and regulated by an unconventional process that remains poorly understood (Alisch et al., 2006).

ORF1p is an RNA binding protein with chaperone activity (Martin and Bushman, 2001, Khazina et al., 2011) while ORF2p contains domains with endonuclease (EN) and reverse transcriptase (RT) activity (Feng et al., 1996, Mathias et al., 1991, Weichenrieder et al., 2004). It is believed that ribonucleoprotein particles (RNPs), consisting of many ORF1p molecules, as few as one or two ORF2ps and one L1 mRNA (Khazina et al., 2011, Basame et al., 2006, Dai et al., 2014) form in the cytoplasm. The RNP is then imported into the nucleus through a still uncharacterized process. L1 RNPs allegedly accumulate in cytoplasmic stress granules (SGs) and processing bodies (PBs) (Goodier et al., 2007, Hu et al., 2015), but the role of these cellular structures in the L1 lifecycle is still controversial. L1 RNPs accumulate mainly in the cytoplasm but nuclear ORF1p was also observed in a certain percentage of cells using cell lines and cancer specimens (Sokolowski et al., 2013, Sharma et al., 2016, Doucet et al., 2016, Goodier et al., 2007, Harris et al., 2010). While several specific antibodies against ORF1p have been raised (Taylor et al., 2013, Wylie et al., 2016, Doucet-O’Hare et al., 2015), a highly effective antibody against ORF2 protein that would allow definitive observation of protein localization is still lacking. Antibodies against LINE-1 ORF2p have been recently developed (De Luca et al., 2016, Sokolowski et al., 2014) but the much lower amount of expressed ORF2p compared to ORF1p makes the study of ORF2p expression and localization difficult. To overcome these difficulties, tagged ORF2ps have been employed (Taylor et al., 2013, Doucet et al., 2010).

In the nucleus, L1 endonuclease nicks the DNA at A/T rich consensus target sites (Feng et al., 1996) and through a process called TPRT (Target Primed Reverse Transcription)(Cost et al., 2002, Luan et al., 1993) inserts a DNA copy into the new genomic target locus. During TPRT, ORF2p EN domain nicks the DNA and the newly formed 3’OH end is then used by the RT domain of ORF2p to prime the synthesis of a complementary DNA using the L1 mRNA as template. A second strand of cDNA is then synthetized and joined to adjacent genomic DNA.

L1 lifecycle is extensively entwined with host cellular processes. Several proteins and cellar pathways have been shown to restrict or support L1 retrotransposition and life cycle (Goodier, 2016). RNA metabolism (Belancio et al., 2008, Dai et al., 2012), DNA damage response (Servant et al., 2017) and autophagy (Guo et al., 2014) are a few cellular processes shown to affect LINE-1 retrotransposition. Progression through the cell cycle was shown by several groups to promote L1 retrotransposition (Shi et al., 2007, Xie et al., 2013) but a molecular understanding of this aspect has been elusive. Because of the importance of the cell cycle in efficient retrotransposition, it has been proposed that, as for some exogenous retroviruses (Goff, 2007, Suzuki and Craigie, 2007), nuclear breakdown during mitosis could represent an opportunity for entrance of L1 into the nucleus (Xie et al., 2013, Shi et al., 2007). This hypothesis, never directly tested previously, was challenged by studies demonstrating effective retrotransposition in non-dividing and terminally differentiated cells (Kubo et al., 2006, Macia et al., 2016).

To shed light on the role of the cell cycle on different aspects of L1 life cycle we explored the nuclear localization of L1 proteins and the retrotransposition efficiency of L1 in different stages of the cell cycle in rapidly dividing cancer cells. Here we use imaging, genetic and biochemical approaches to show that in these cells, the L1 lifecycle is intimately coordinated with the cell cycle. LINE-1 encoded proteins enter the nucleus during mitosis and retrotransposition appears to occur mainly during S phase.

## RESULTS

### Analysis and quantification of ORF1p and ORF2p expression and cellular localization

Previous works (Doucet et al., 2016, Luo et al., 2016, Sokolowski et al., 2013, Goodier et al., 2007, Goodier et al., 2004, Sharma et al., 2016, Rodic et al., 2014, Taylor et al., 2013, Branciforte and Martin, 1994) have shown varying localization of ORF1 protein in cells growing in culture or in mammalian specimens from various organs and tumors. In particular, the nuclear localization of LINE-1 encoded proteins has been sparsely studied and the mechanisms driving import of L1 retrotransposition intermediates into the nucleus are largely unknown.

We therefore set out to characterize the cellular localization of L1 ORF1p and ORF2p in human cells overexpressing recoded (ORFeus) or non-recoded L1 (L1rp) with a 3xFlag tag on the ORF2p C-terminus (Taylor et al., 2013). Immuno-fluorescence staining of HeLa-M2 cells (Hampf and Gossen, 2007) overexpressing ORFeus, showed clear expression of ORF1p and ORF2p (Fig. 1A and D and S2-5). As previously observed (Taylor et al., 2013), ORFeus ORF1p was expressed in virtually all the cells (@ 97%) whereas ORF2p, encoded by the same bicistronic L1 mRNA also expressing ORF1p, was expressed in just a subset of cells (@10% using a rabbit anti-Flag antibody and @20% using a more sensitive mouse anti-Flag antibody) (Fig. 1B, bar graph and inset). This pattern of expression is most likely due to an unknown mechanism controlling ORF2p translation (Alisch et al., 2006). Interestingly, when non-recoded L1 was over-expressed, only 44% of cells displayed ORF1p expression. Overexpression of ORFeus-Hs and L1rp with or without the L1 5’ untranslated region (5’ UTR) excluded the possibility that the reduced expression of L1rp compared to ORFeus was due to the presence of the 5’ UTR (Fig. S1) (Chen et al., 2012). Due to the overall lower expression of L1rp, ORF2p was barely observable in cells expressing non-recoded L1 (Fig. 1A and B). ORF1p was mainly present in the cytoplasm but some cells (@26%) also showed clear nuclear staining (Fig. 1A, Fig. 1D, white arrowheads, Fig. S2-5). Interestingly, the cells with nuclear ORF1p fluorescence were usually observed as pairs of cells in close proximity. This observation suggested to us that cells with nuclear ORF1p may have undergone mitotic division immediately prior to fixation and observation. In HeLa cells, this observation was made more evident by the fact that daughter cells displaying nuclear ORF1p were sometimes connected by intercellular bridges often containing filaments of DNA and persisting from incomplete cytokinesis during the previous mitosis (Fig. 1D, lower panels) (Steigemann et al., 2009, Carlton et al., 2012). To quantify our initial qualitative observation of closer cell proximity for cells displaying nuclear ORF1p, we employed automated image acquisition of HeLa cells expressing recoded L1 and stained for ORF1p (see “quantification and statistical analysis” section). We compared the distance of pairs of cells expressing nuclear ORF1p versus the distance of the same number of cell pairs expressing ORF1p just in the cytoplasm. This “proximity analysis”, described in more details in the methods section, clearly shows that cells with nuclear ORF1p are statistically closer to each other than cells expressing ORF1p exclusively in the cytoplasm (Fig. 1C).

**Fig. 1.**
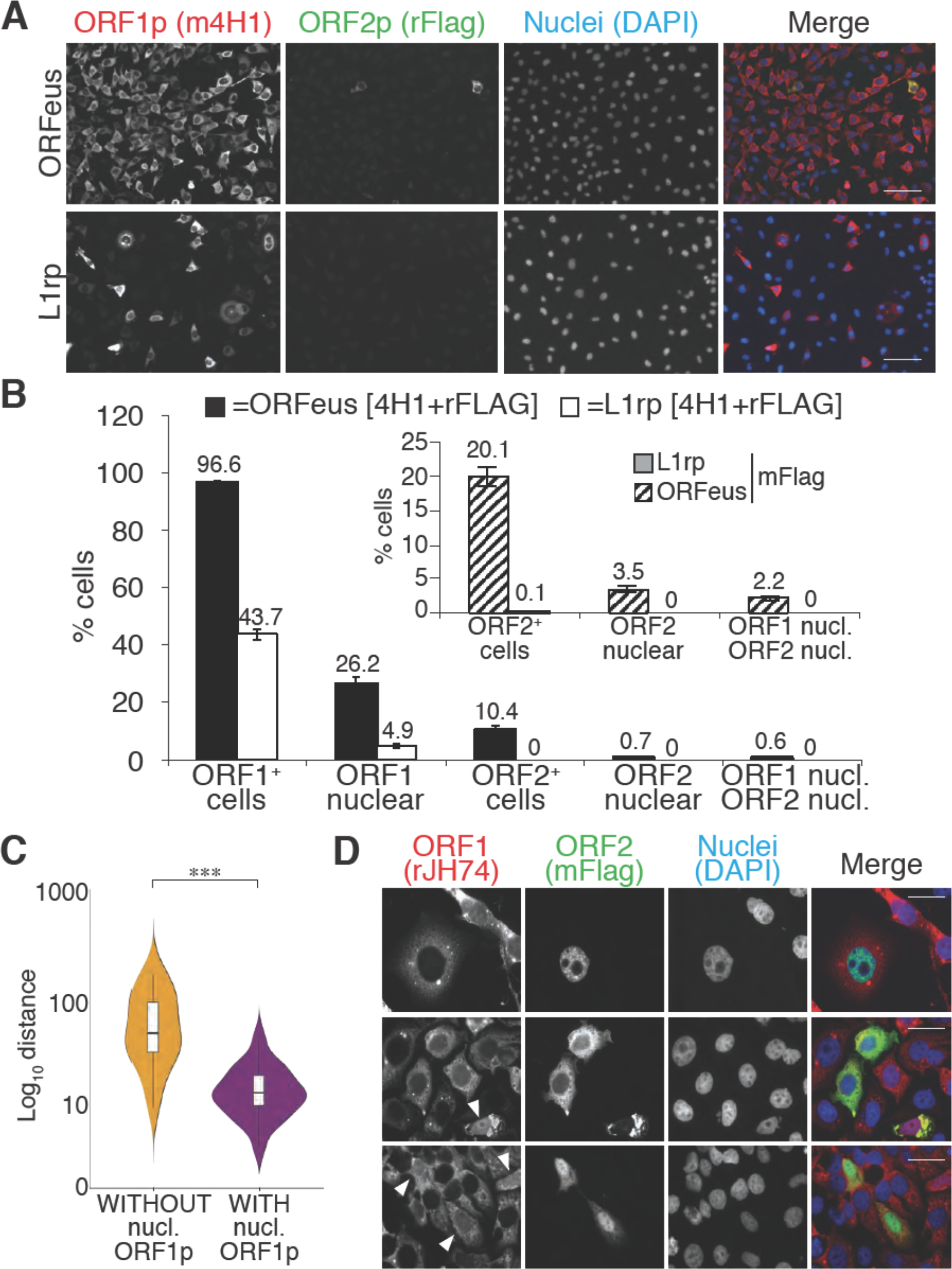
L1 proteins expression and localization. (A) Representative pictures of immunostained HeLa M2 cells expressing a recoded L1 (ORFeus) or a non-recoded L1 (L1rp). Antibody target names are reported atop the corresponding pictures and colored according to the colors used in the merged pictures. Scale bar= 100µm. (B) quantification of ORF1p and ORF2p expression in the cytoplasm and nucleus of HeLa cells expressing the indicated L1 element (error= S.E.M.) of at least 10 20X fields with about 100 cells each. The inset shows quantification of ORF2p stained using an anti-FLAG mouse antibody (SIGMA-M2 F1804). (C) Proximity analysis of cells with (yellow) or without (purple) nuclear ORF1p is reported. HeLa cells expressing Flag tagged ORFeus for 24 hours, were stained with anti-ORF1p JH74 Ab and DAPI. Pictures were collected using an Arrayscan microscope and analysis was performed as described in the Experimental Procedures. ***=p <0.001 (p= 4.351e^-13^) (D) Immunostaining of ORF1p and ORF2p using, respectively, rabbit monoclonal Ab JH74 and mouse monoclonal antibody M2 against FLAG. Scale bar= 20µm. White arrowheads indicate cells with nuclear ORF1p.

The cytoplasmic ORF1p localization pattern was often dominated by previously described cytoplasmic foci (Doucet et al., 2010, Goodier et al., 2007). The formation of these foci is particularly enhanced by L1 overexpression. To avoid formation of such structures, which are presumed artifacts induced by non-physiological expression levels of ORF1p, we therefore performed the localization experiments reported here using Tet-inducible constructs, and we induced L1 expression at lower concentrations of doxycycline (0.1µg/ml) compared to the concentrations typically used to induce full induction of retrotransposition or production of RNP intermediates for proteomic studies (1 µg/ml; (Taylor et al., 2013). We observed very heterogeneous localization of ORF1p and ORF2p in cells overexpressing L1 upon treatment with 0.1 µg/ml doxycycline for 24 hours. ORF1p is observed to be only cytoplasmic (Fig. 1D top panel), or both cytoplasmic and nuclear (Fig. 1D, top panel but is never exclusively nuclear when immunostaining is performed using 4H1, 4632 and JH74 antibodies (Fig. S2-6). ORF2p also had heterogeneous nuclear/cytoplasmic localization with cells displaying ORF2p only in the cytoplasm (Fig. 1A and Fig. 1D middle panel), only in the nucleus (Fig.1D, top panel) or in both the cytoplasm and the nucleus (Fig. 1D, lower panel). Nuclear ORF2p had a punctate pattern (Fig. 1D top panel) whereas nuclear ORF1p was uniformly distributed in the nucleus and usually excluded from nucleoli (Fig.1D, arrowheads). Also, a small population of cells (usually less than 0.1% of L1 expressing cells) showed a strong and clear nuclear signal of ORF2p in the absence of nuclear ORF1p (Fig.1D, top panel and S7) consistent with a possible nuclear form of L1 RNPs that lacks ORF1p (see also accompanying paper by Molloy et al.). On the contrary, in the cytoplasm, ORF2p always co-localized with ORF1p (Fig. 1A and D).

We compared several antibodies against ORF1p that confirmed a consistent pattern of nuclear/cytoplasmic localization of ORF1p (Fig. S2). The mouse 4H1 monoclonal antibody (mAb) recognizing the N-terminus of ORF1p and the rabbit JH74 antibody against the C-terminus of ORF1p have an identical pattern. The polyclonal 4632 antibody (pAb) displayed a high nuclear non-specific signal that renders this antibody less sensitive than 4H1 and JH74 (Fig. S2). The JH73g rabbit Ab was distinct from the other antibodies and recognized mainly nuclear ORF1p (Fig. S2 A and B and S6). This nuclear form of ORF1p is also recognized by 4H1 and JH74 antibodies but is much more easily observed using JH73g because of its higher affinity for nuclear ORF1p and much lower affinity for cytoplasmic ORF1p. Indeed, quantification of nuclear ORF1p upon staining with JH74 and JH73g revealed a comparable percentage of cells displaying nuclear ORF1 using the two antibodies (@14-17%). The percentage of cells detected overall as expressing ORF1p using JH73g Ab is much lower than the percentage recognized as ORF1p expressing cells by JH74 Ab (37% versus 93% respectively) because JH73g is able to recognize mainly the nuclear form (Fig. S2B). This unusual staining pattern suggests that the nuclear form of ORF1 may be highly enriched for a conformation specifically recognized by JH73g. Staining of L1rp expressing cells, on the other hand, reveals a lower threshold of sensitivity for the JH73g antibody that is unable to detect the lower amount of non-recoded ORF1p. Interestingly, ORF1p immunoprecipitated with JH73g Ab is impaired in binding to ORF2p protein, consistent with the possibility that most of the nuclear ORF1p species fail to bind ORF2p (Fig. S2D).

### ORF1p enters the nucleus during mitosis

Our immunofluorescence staining of ORF1p and ORF2p suggest that L1 RNPs enter the nucleus during mitosis, when the nuclear membrane breaks down. To better explore this hypothesis, previously put forward by other studies (Xie et al., 2013), we exploited well-characterized markers of the cell cycle: geminin, expressed only in S/G2/M phases, and Cdt1, which specifically marks the G1 phase (Arias and Walter, 2007). Co-staining of ORF1p with geminin and Cdt1 clearly showed that ORF1p is nuclear in cells expressing Cdt1 (G1 phase), and completely cytoplasmic in cells expressing geminin (Fig. 2A and quantification in B). We also verified these results using the FUCCI system (Fluorescent Ubiquitin Cell Cycle Indicator), that exploits Cdt1 and geminin fragments fused to mAG and mKO2 (Monomeric Azami-Green and Monomeric Kusabira-Orange 2 fluorescent proteins) respectively (Sakaue-Sawano et al., 2008). Expression of an ORF1p-HaloTag7 in HeLa.S-FUCCI cells treated with the Halo tag ligand JF646, (Grimm et al., 2015) shows that ORF1p is nuclear only in cells in G1 phase with orange nuclei resulting from expression of mKO2-tagged Cdt1 fragment (Fig. 2C). In this setting, no instances of cells with nuclear ORF1p-Halo and green nucleus (cells in S/G2/M) were observed. ORF2p-Halo was expressed in cells scattered throughout the cell cycle that displayed either orange (Cdt1 expressing) or green (geminin expressing) nuclei (Fig. S7). This result suggests that ORF2p expression is not cell cycle regulated. Overall, our analyses strongly suggest that ORF1p, most likely together with ORF2p, enters into the nucleus during mitosis and is retained there when the nuclear membrane reforms after cell division.

**Fig. 2.**
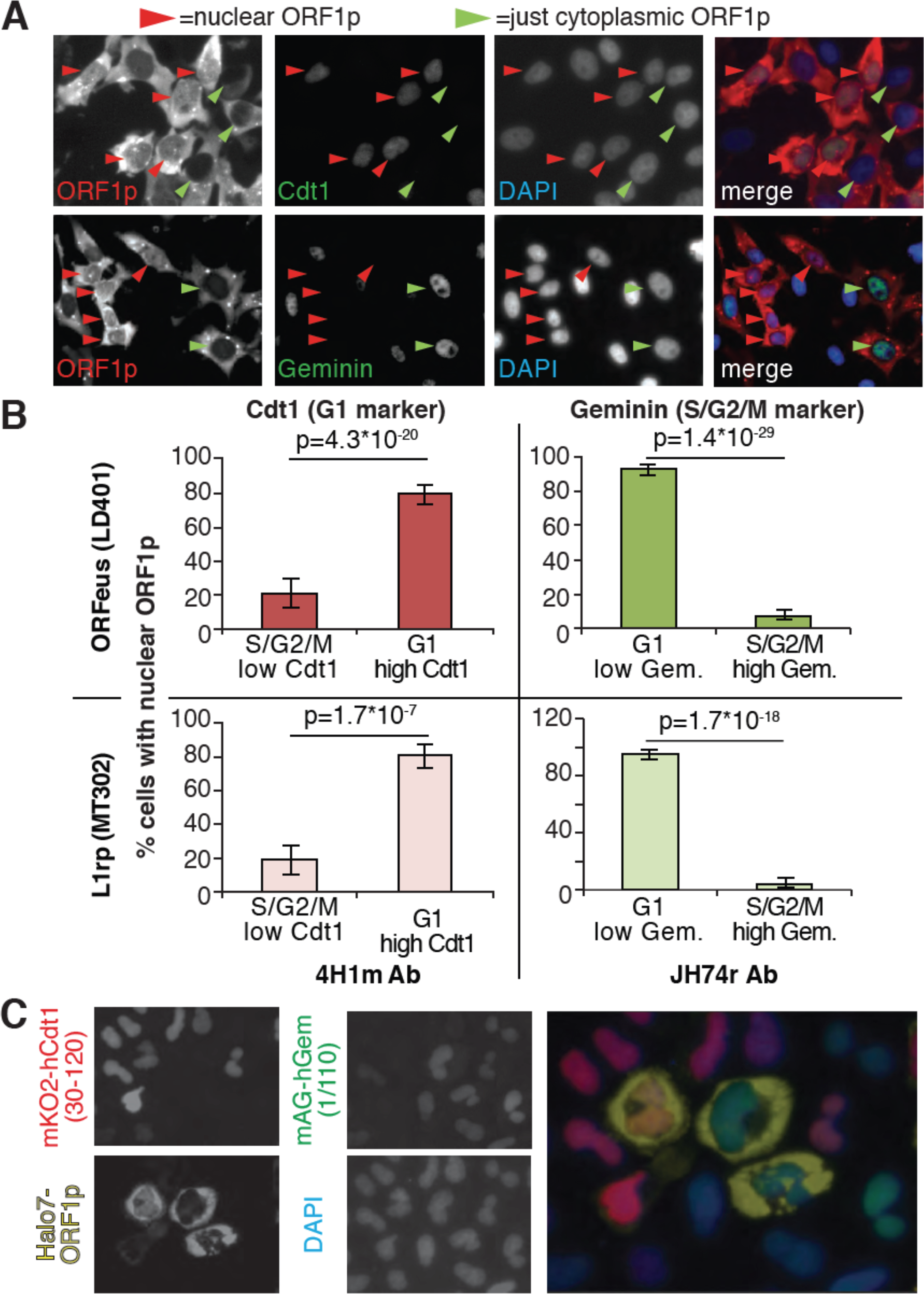
ORF1p localizes in the nucleus immediately after mitosis. (A) Representative pictures of co-immunostaining of ORF1p with Cdt1 (marker of G1 phase) or geminin (marker of S/G2/M phase). DNA was stained with DAPI. Red arrowheads= cells with nuclear ORF1p, green arrowheads= cells without nuclear ORF1p. (B) quantification of cells expressing nuclear ORF1p and Cdt1 (red, left top and bottom) or Geminin (green, right top or bottom). HeLa M2 cells expressing ORFeus (plasmid LD401) or L1rp (plasmid MT302) are shown in the two top or bottom panels respectively. The antibodies used for staining and quantification are indicated at the bottom of the graphs. (error=S.E.M.). (C) Representative pictures of live HeLa. S-FUCCI cell lines expressing rtTA and an inducible recoded L1 with ORF1p C-terminally tagged with an Halotag7 (Ohana et al., 2009). JF646 ligand (Grimm et al., 2015) was used to visualize ORF1-Halotag7. The proteins visualized are reported on top of the pictures in colors used in the merged picture (bottom picture).

A direct consequence of these conclusions would be that prolonged expression of L1 in dividing cells should eventually lead to a population with all cells displaying nuclear ORF1p because a longer time of L1 induction will allow all the cells to undergo mitosis while expressing ORF1p. To test this hypothesis, we quantified the percentage of cells displaying nuclear ORF1p after 24 and 48 hours of L1 expression, considering that HeLa cell doubling time is about 24 hours. Automated picture collection (Arrayscan HCS, Cellomics) and software based nuclear/cytoplasmic analysis (HCS studio cell analysis software) was implemented as for proximity analysis. We set very stringent negative fluorescence thresholds (limit in Fig. 3A) determined from cells not treated with doxycycline and therefore not expressing L1, as described in the methods section. These stringent parameters were necessary to avoid interference of the strong ORF1p cytoplasmic signal with the measurement of nuclear ORF1p signal. The analysis of ORF1p nuclear and cytoplasmic distribution surprisingly showed that the percentage of cells with nuclear ORF1p does not increase but actually decreases after 48 h of L1 induction compared to the 24 h time point (Fig. 3A). The decrease in nuclear ORF1p after 48 h induction is probably due to the decreased growth rate of a more confluent cell population. The absence of an increase of cells with nuclear ORF1p with increased time of L1 induction, suggests that, after entering the nucleus in M phase, nuclear ORF1p is either degraded and/or exported from the nucleus during or after G1 phase.

**Fig. 3.**
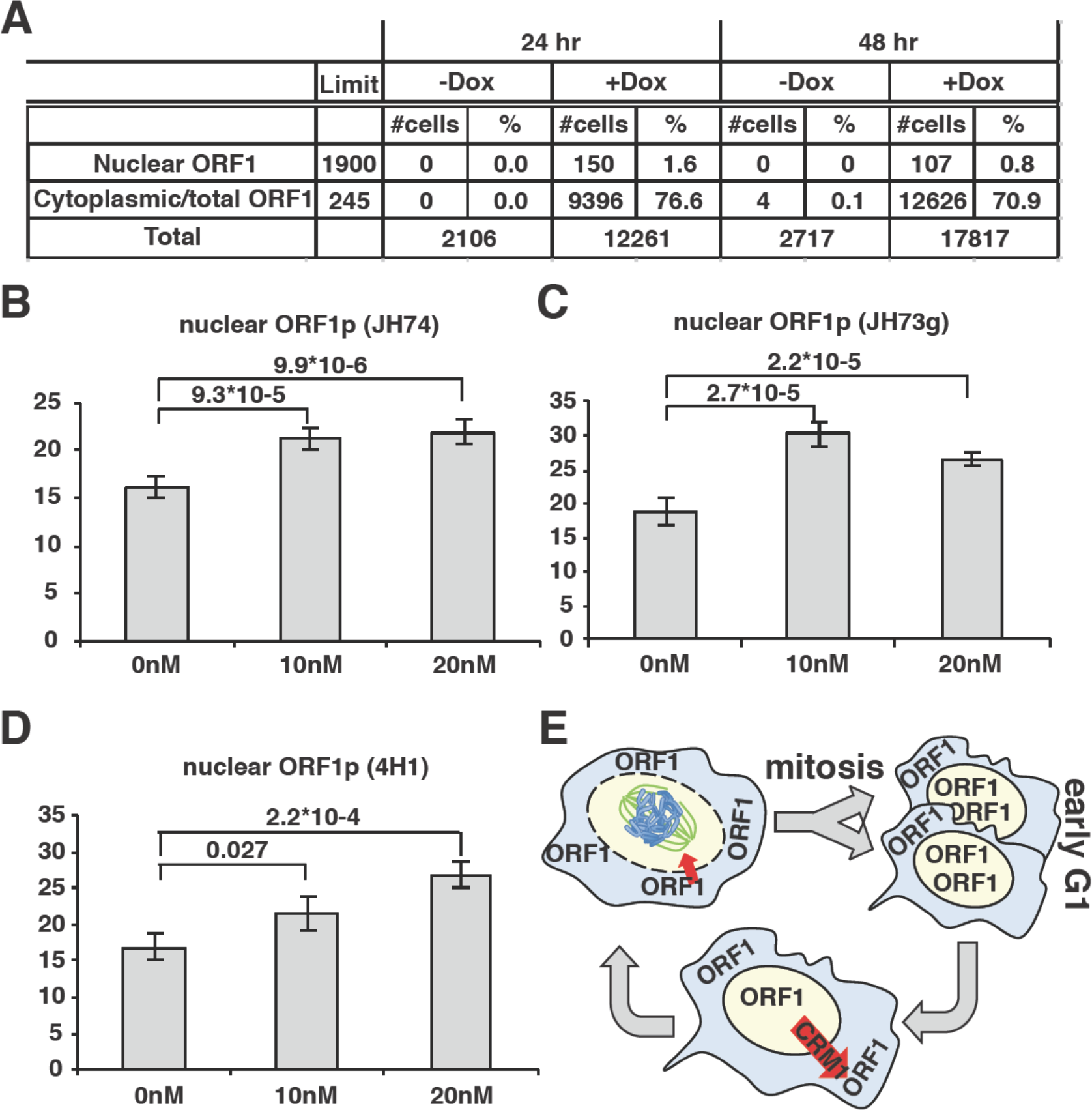
ORF1p nuclear localization upon leptomycin treatment. (A) HeLa M2 cells expressing a recoded L1 (ORFeus) with Flag tagged ORF2p were treated for 24 or 48h with 0.1µg/ml doxycycline on chamber slides. After treatment, cells were fixed in formalin and stained with JH74 primary antibody, Alexa 647 labeled secondary antibody and DAPI. Slides were scanned with Arrayscan and analyzed using Image Studio HCS software. The number of cells with cytoplasmic ORF1p (also considered as the total number of ORF1p expressing cells) and nuclear ORF1p are reported as well as total amount of cells calculated from DAPI staining. (B-D) quantification of cells expressing nuclear ORF1p in HeLa M2 cells treated with or without leptomycin 10 or 20nM for 15 hrs. After treatment cells were fixed and ORF1p stained using the indicated antibodies. Two tailed T-test p-values are reported for significant differences (error =S.E.M.) (E) Schematic of ORF1p nuclear/cytoplasmic dynamics during the cell cycle.

### ORF1p nuclear localization is increased upon leptomycin treatment

To better explore potential cytoplasmic/nuclear shuttling of ORF1p and ORF2p we took advantage of a known inhibitor of exportin 1 (XPO1/CRM1), leptomycin b. We treated HeLa cells expressing LINE-1 with leptomycin for 18 h. Two different concentrations of leptomycin were used and several antibodies (Abs) were utilized to detect ORF1p in immunofluorescence assays (Fig. 3B-E). At both leptomycin concentrations, and using any of the Abs recognizing ORF1p we observed an increased number of cells with nuclear ORF1p after leptomycin treatment, suggesting that at least a subset of ORF1p is exported from the nucleus in a CRM1-dependent manner (Fig. 3E).

### LINE-1 retrotransposition peaks during S phase

Our results suggest that ORF1 protein, presumably in a ribonucleoprotein complex with ORF2p and L1 mRNA, is able to enter the nucleus during mitosis and it accumulates in the nucleus in early G1 phase of the cell cycle. Following early G1, ORF1p is then exported to the cytoplasm through a CRM1 dependent mechanism. We therefore asked whether L1 retrotransposition occurred in a cell cycle-dependent manner and more specifically during M phase, when we observed ORF1p in the nucleus and when chromatin is accessible to L1 RNPs. To answer this question we performed retrotransposition assays using a previously described ORFeus-GFP-AI reporter (Taylor et al., 2013, An et al., 2011). HeLa cells expressing the retrotransposition reporter were treated for increasing times with nocodazole (Fig. 4A), a cell cycle inhibitor that blocks cells in M phase interfering with microtubule assembly (Ma and Poon, 2017, Rosner et al., 2013). Increasing time of nocodazole treatment, and therefore longer times in M phase, fails to increase the percentage of M phase green cells (Fig. 4A-B), suggesting that L1 retrotransposition does not occur during M phase. Longer times of nocodazole treatment (21hrs) increased cell death, detected by an increase of positive propidium iodide-positive cells, and a consequent decrease in retrotransposition (Fig. 4B, dotted line).

**Fig. 4.**
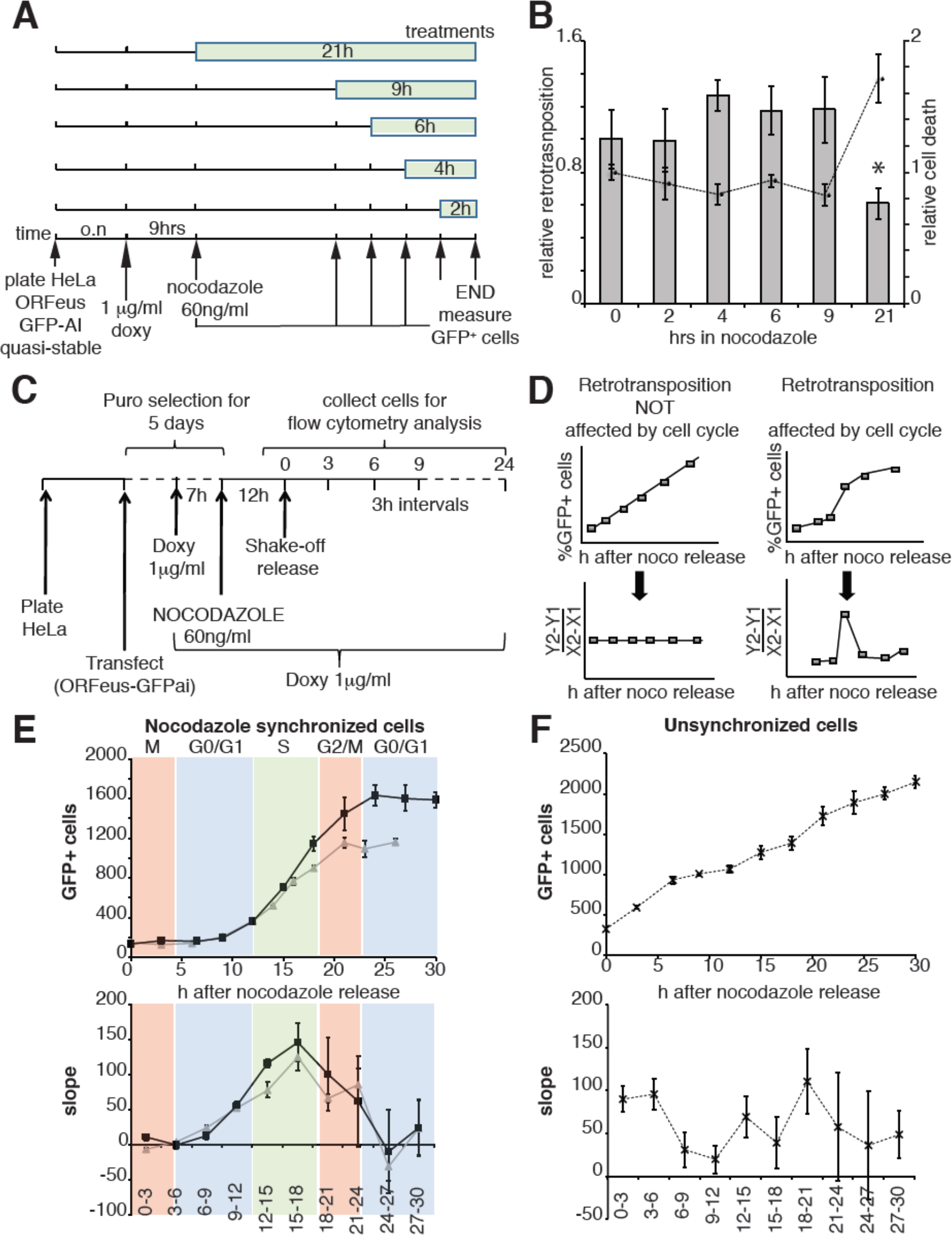
Retrotransposition during the cell cycle. (A) Scheme of the experimental timeline followed for experiments presented in b). HeLa M2 cells stably expressing episomal ORFeus-GFP-AI reporter (EA79 plasmid) were plated in 10cm tissue culture treated plates. After overnight incubation expression of L1 was induced with 1 µg/ml doxycycline. After 9 h induction, nocodazole was directly added to the media at various times in different plates to a final concentration of 60 ng/ml. After 30 h from the beginning of doxycycline treatment all cells were collected and the percentage of GFP^+^ cells measured using a flow cytometer as described in the method section. (B) Histogram boxes (left Y axes) represent the relative % of GFP^+^ cells measured by cytofluorometry after treatments described in a) The dots connected by a dotted line (right Y axes) represent the relative % of PI^+^ cells (dead cells). Retrotransposition and PI percentages of cells not treated with nocodazole (time 0) was set as 1. (error=S.D., *= p<0.05. (C) Scheme of the experimental timeline followed for experiments presented in E-F). HeLa M2 cells were plated and transfected in 6 well plates. 24 h after transfection 1µg/ml puromycin was added to the medium for 5 days. During puromycin selection cells were split into 10 cm plates to avoid contact inhibition. During the 5^th^ day of puromycin selection, 3x10^6^ cells were freshly replated in 10 cm plates and after 3 h, doxycycline was added to the media to a final concentration of 1µg/ml. After 7 h, nocodazole was added to a final concentration of 60 ng/ml. After 12 hours, media was discarded and mitotic cells were collected by mechanical shake off. Cells were washed and 0.4x10^6^ mitotic cells were replated in each of 3 cm wells. At the indicated time points the percentage of GFP^+^ cells was measured using a flow cytometer. (D) prediction of retrotransposition measurements if cell cycle affects (left panels) or does not affect (right panels) retrotransposition. (E-F) Retrotransposition analysis of nocodazole synchronized (E) or not synchronized (F) cells. The % of GFP^+^ cells in a population of 10000 cells is reported in the indicated time points. Bottom panels show slopes changes from the corresponding measurements on the top panels. The shaded colored boxes indicate specific cell cycle stages extrapolated from propidium iodide (PI) measurements reported in Fig. S8. The two lines (gray and black) represent two experiments using independent transfections of the retrotransposition reporter. (error=S.D., n=4).

We then expanded our analysis, measuring retrotransposition during a single cell cycle in a population of HeLa cells synchronized by nocodazole treatment, subsequent “mitotic shake off” and released into the cell cycle in the absence of nocodazole (Fig. 4A). Measurements of the percent of cells that underwent retrotransposition were performed every three hours starting after release from nocodazole synchronization. The cell cycle stage of the cells at each time point was determined by propidium iodide staining (Fig. S8C). A linear increase of retrotransposition should be observed if retrotransposition is unbiased towards specific cell cycle stages, while a non-linear increase represents a specific stage at which retrotransposition is enhanced. Calculation of the slope of the increase of GFP^+^ cells should therefore produce a clear peak at the time during which most retrotransposition occurs (Fig. 4D). This approach allowed us to identify a peak of retrotransposition in the S phase (Fig. 4E top and bottom left panels). Control non-synchronized cells, as expected, showed a linear increase in retrotransposition and no clear peaks were identified (Fig. 4F top, bottom right panels).

To evaluate whether the cell cycle controls retrotransposition using a method independent of cell synchronization, we developed a fluorescent-AI reporter that introduces a temporal component to canonical retrotransposition reporters. To this end we utilized the previously characterized monomeric fluorescent timers (FT) (Subach et al., 2009). These derivatives of mCherry change their fluorescence emission from blue to red over 2 to 3 hours. We introduced an antisense intron within the coding region of the “fast-FT” and inserted this cassette in the 3’UTR of a recoded L1 (Fig. 5A). Transfection of the L1-fastFT-AI construct into HeLa cells allowed us to identify cells that underwent retrotransposition within a ∼3 hr period preceding the analysis, as reported by previous work (Subach et al., 2009). Our quantification also supports the previously reported timing of FT maturation (Subach et al., 2009) with an average conversion time from blue to red of 2.35±0.52 hours (Fig. S8B and Fig. 5A). Immediately after L1-fastFT-AI retrotransposition, the fast FT is expressed and the cells emit blue fluorescence. Quantification of GFP expression upon doxycycline induction revealed that 50% of the cells expressed visible GFP at 2.71±0.46 hours and 90% of the cells expressed visible GFP within 8.01±0.63 hours of doxycycline treatment (Fig. S8A and Fig. 5A). Upon maturation (2.35±0.52 hours) the blue proteins begin turning red. The cells are now marked by a blue population of proteins continuously transcribed by the constitutive CMV promoter and a red population of aged proteins matured from the blue form (Fig. 5A). Analysis of the cell cycle stage of FACS sorted “blue only” cells (cells that underwent retrotransposition within 3 h of the analysis) compared to fluorescence negative cells FACS sorted from the same population (cells that did not undergo retrotransposition before analysis) revealed a strong enrichment in S phase cells, a partial enrichment in cells in the G2/M phase and a strong de-enrichment of cells in G1 phase (Fig. 5B and Fig. S9 A-B). These results confirmed the strong bias of retrotransposition towards S phase that we measured using nocodazole synchronization (Fig. 4).

**Figure 5.**
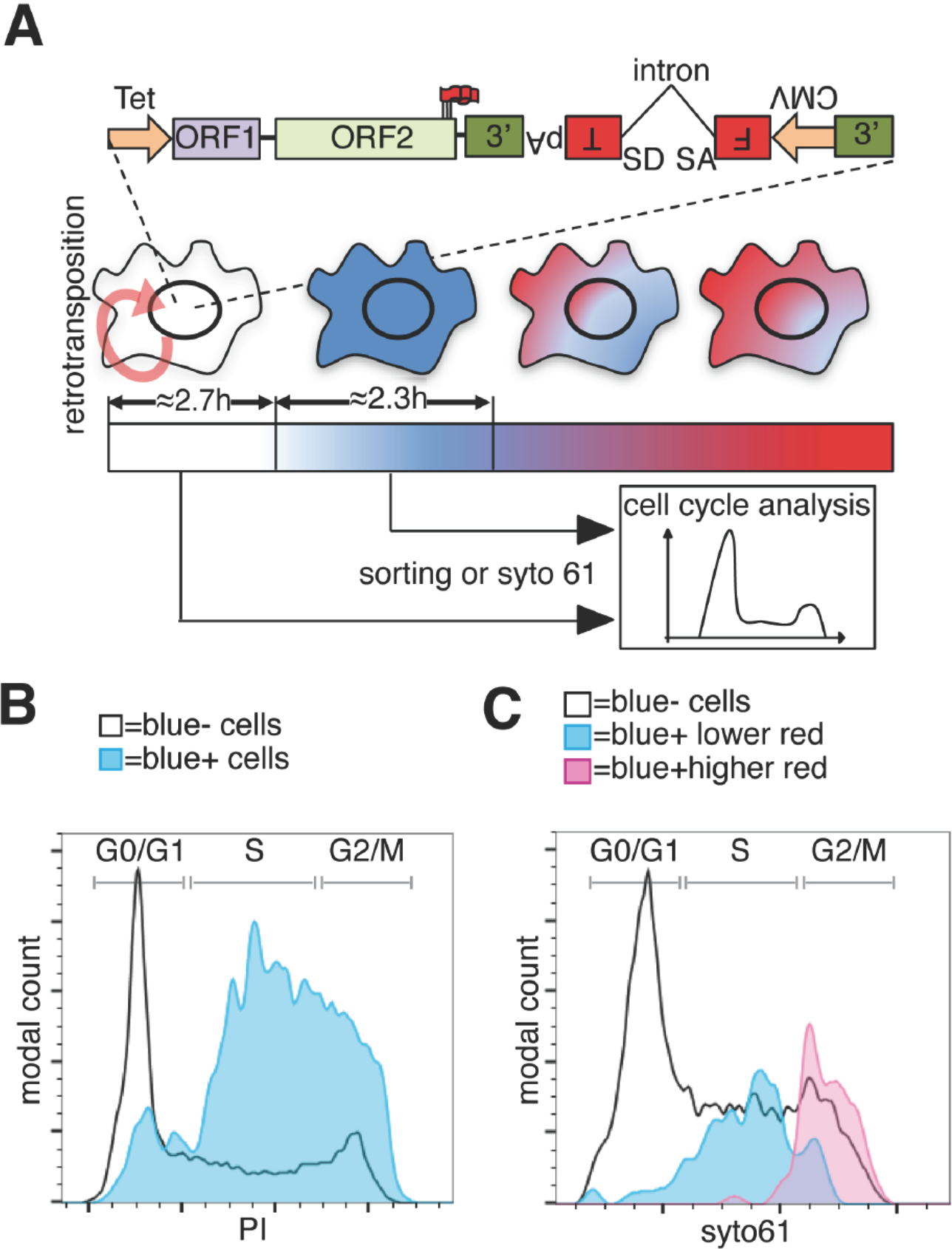
Analysis of L1 retrotransposition during the cell cycle using FT-AI reporter. (A) Schematic of the fluorescent-timer-AI reporter (FT-AI) and of the experimental design. HeLa M2 cells were transfected with a FT-AI reporter. After induction of L1 expression (24 h, 1µg/ml doxycycline), cells undergoing retrotransposition are blue and can be sorted by FACS or directly stained with syto61 for cell cycle analysis. Within about 2.7 h of doxycycline treatment 50% of the cells will start to express the FT reporter as estimated in Fig. S8A. After about 2.35 h (Fig. S8B) from expression of the blue FT (retrotransposition event), cells start to become red. Double negative (blue-/red-) cells were also collected by sorting and analyzed as control. The sorted cells are then stained with PI and their cell cycle stage determined. (tet=Tetracycline inducible promoter, 3’= 3’UTR, pA=polyA signal, FT=fluorescent timer, SD=splice donor, SA=splice acceptor, CMV= cytomegalovirus constitutive promoter, red round arrow=L1 “jumping”). (B) Histogram of cell cycle distribution of sorted and PI stained blue^-^red^-^ cells (black line) and blue^+^red^-^ cells (blue histogram). The percentage of cells in each cell cycle stage after sorting and PI staining are reported in Fig. S9. (C) Cell cycle analysis using syto61 dye of blue^-^ cells that did not undergo retrotransposition (black line), of blue^+^ cells expressing lower red signal (blue histogram) or blue^+^ cells expressing higher red signal (purple histogram). The complete profiles of analysis for the reported cells are presented in Fig. S9.

To better dissect the cell cycle stage of cells that underwent retrotransposition, we also implemented a second approach that does not involve cell sorting but simply allows direct analysis of the cell cycle in cells expressing the FT-AI reporter (Fig. 5C and Fig. S9). After 24 h of doxycycline treatment, cells expressing the L1-fastFT-AI reporter were directly stained with syto61 DNA labeling dye and analyzed. The main population of blue negative cells that did not undergo retrotransposition, showed cells distributed throughout the cell cycle (G1=49.5%, S=34.2%, G2/M=15.2%) (Fig. 5C, black line and Fig. S9C-D). Using this analysis we were able to divide the population of blue positive cells (blue^+^) in two subpopulations: cells with relatively higher red fluorescence (Fig. 5C, purple profile and Fig. S9C), and cells with lower red fluorescence (Fig. 5C, blue profile and Fig. S9C). The former group of cells underwent retrotransposition in a time closer to the time of analysis compared to the blue^+^ cells with higher red fluorescence. Consistent with the previous experiments, the blue^+^ cells with lower red fluorescence (blue peak) are mainly in S phase (G1=9.38%, S=78.1%, G2/M=12.5%). Blue^+^ cells with higher red fluorescence (purple peak), which had more time to proceed through the cell cycle after retrotransposition and before analysis (from S to G2/M), were mainly in G2/M phase (G1=0%, S=10.9%, G2/M=89.1%). This result clearly shows that the wide peak of “blue only” sorted cells that spread across S and G2/M phases (Fig. 5B) actually consists of two subpopulations/peaks: a population of cells in S phase that underwent retrotransposition a short time before analysis and a second population of cells in G2/M phase that underwent retrotransposition earlier relative to analysis.

These observations collectively indicate that L1 retrotransposition has a strong cell cycle bias and preferentially occurs during the S phase.

### ORF2p binds chromatin and localizes at replication forks with PCNA during S phase

To gain biochemical insight into the timing of L1 retrotransposition we investigated the timing with which ORF2p was recruited onto chromatin, a necessary step for retrotransposition. We isolated nuclear soluble and chromatin bound proteins from cells synchronized and released into the cell cycle, as in Fig. 4C. Immunoblot analysis showed no differences in the amount of histone H3 and ORF1p present on chromatin and, as expected, the analysis revealed chromatin recruitment that peaked in S phase for PCNA (Strzalka and Ziemienowicz, 2011) and Upf1 (Azzalin and Lingner, 2006). Supporting our previous results, ORF2p was recruited on chromatin in S phase in a similar manner to Upf1 and PCNA (Fig. 6A-B).

**Fig. 6.**
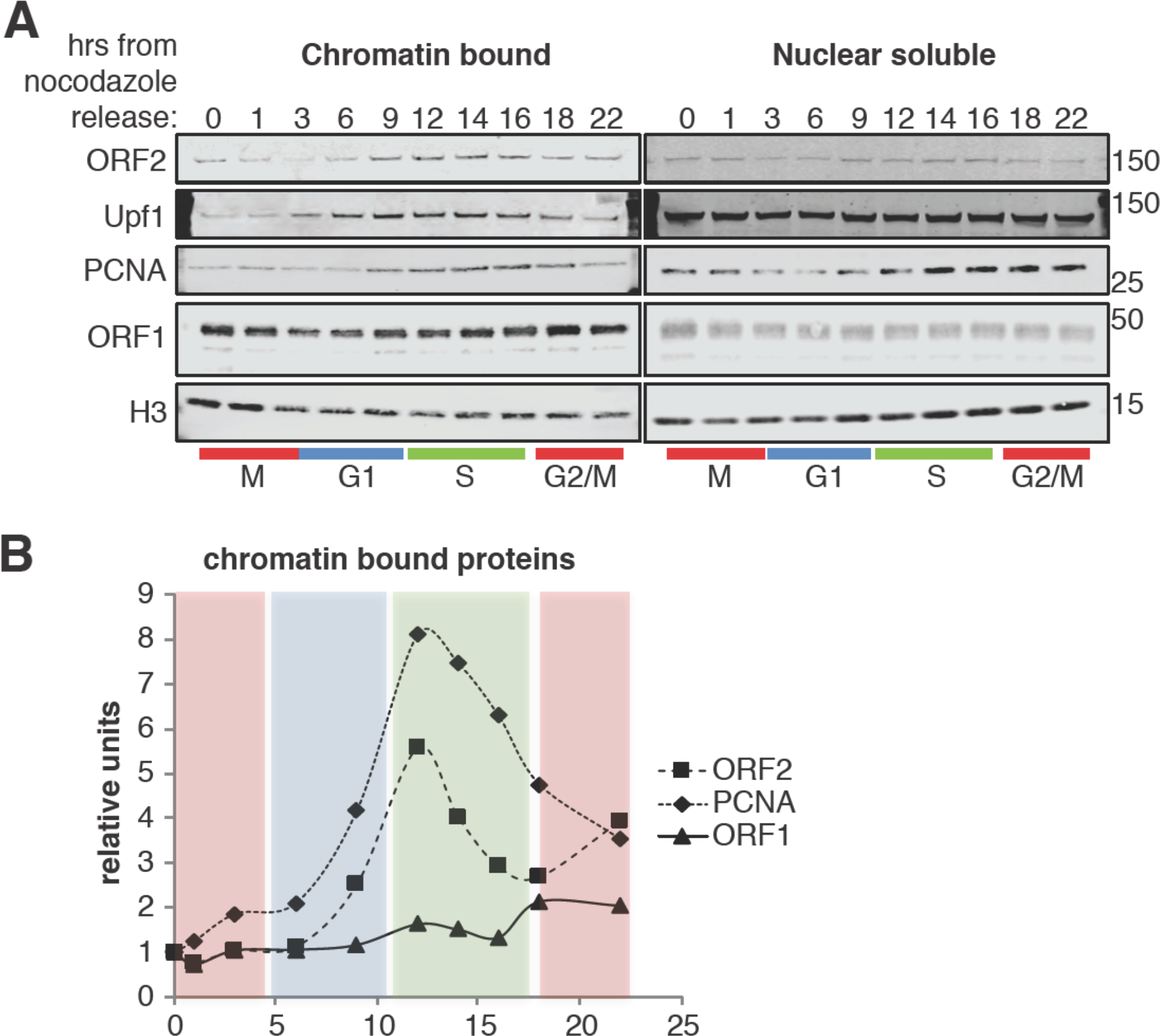
ORF2p binds DNA during S phase. (A) Western blot of chromatin bound proteins or nuclear soluble proteins extracted from HeLa M2 cells expressing recoded L1, synchronized with nocodazole and released into the cell cycle for the indicated times. The colored bars below the blots indicate cell cycle phases extrapolated from Fig. S8. The targets of the antibodies used for blotting are indicated. The quantification of the ORF2p, ORF1p and PCNA proteins bound to chromatin are reported in (B) as ratio of H3 signals. The relative units at time zero are set to 1.

We previously showed that ORF2p binds PCNA through a PIP domain in the ORF2 protein, sandwiched between the EN and RT domains, and that the PCNA-ORF2p interaction is necessary for retrotransposition in HeLa and HEK293 cells (Taylor et al., 2013). The PCNA-ORF2p complex is mainly chromatin bound (Fig. S10), supporting the idea that PCNA binds ORF2p during retrotransposition. We therefore followed up on these previous findings investigating the interactome of the ORF2p-PCNA complex specifically. To this end we engineered a V5-tag at the N-terminus of PCNA in HCT116 cells stably expressing a doxycycline inducible ORFeus. We performed sequential immunoprecipitation of ORF2p followed by V5/PCNA IP (Fig. 7A) and we analyzed the interacting partners of the ORF2p-PCNA complex by mass spectrometry. We identified several MCM proteins (MCM3, MCM5 and MCM6) as well as TOP1 (DNA topoisomerase 1), PARP1 (Poly [ADP-ribose] polymerase 1) and RPA1 (Replication Protein A1) (Fig. 7B). These proteins are known to be co-recruited with PCNA on the origins of DNA replication before S phase (MCM proteins) and during S phase on the replication fork (MCM, PCNA, TOP1, RPA1 and PARP1 proteins) (Czubaty et al., 2005, Remus et al., 2009, Ying et al., 2016). Co-immunoprecipitation of Flag/ORF2 or ORF1 proteins from HEK293 cells expressing ORFeus, recapitulated the interaction of ORF2p with MCM6 and PCNA (Fig. 7B). As expected, immunoprecipitation of ORF2p pulled down a fraction of ORF1p, but also MCM6 and PCNA proteins. Interestingly, as expected from our previous observations revealing that ORF1p is not necessarily in the complex(es) with chromatin bound retrotransposing L1 RNPs, immunoprecipitation of ORF1p pulled down only a small amount of ORF2p, and also a smaller amount of MCM6 and PCNA proteins. These observations suggest that the retrotransposing L1 complex containing ORF2p, PCNA and components of the replication fork such as MCM6, are depleted of ORF1 proteins. Finally, we performed immunofluorescent staining of ORF2p and PCNA in HeLa cells synchronized in S phase by double thymidine synchronization. A subset of nuclear ORF2p puncta overlapped with PCNA foci, marking potential regions of active DNA replication (Fig. 7D).

**Fig. 7.**
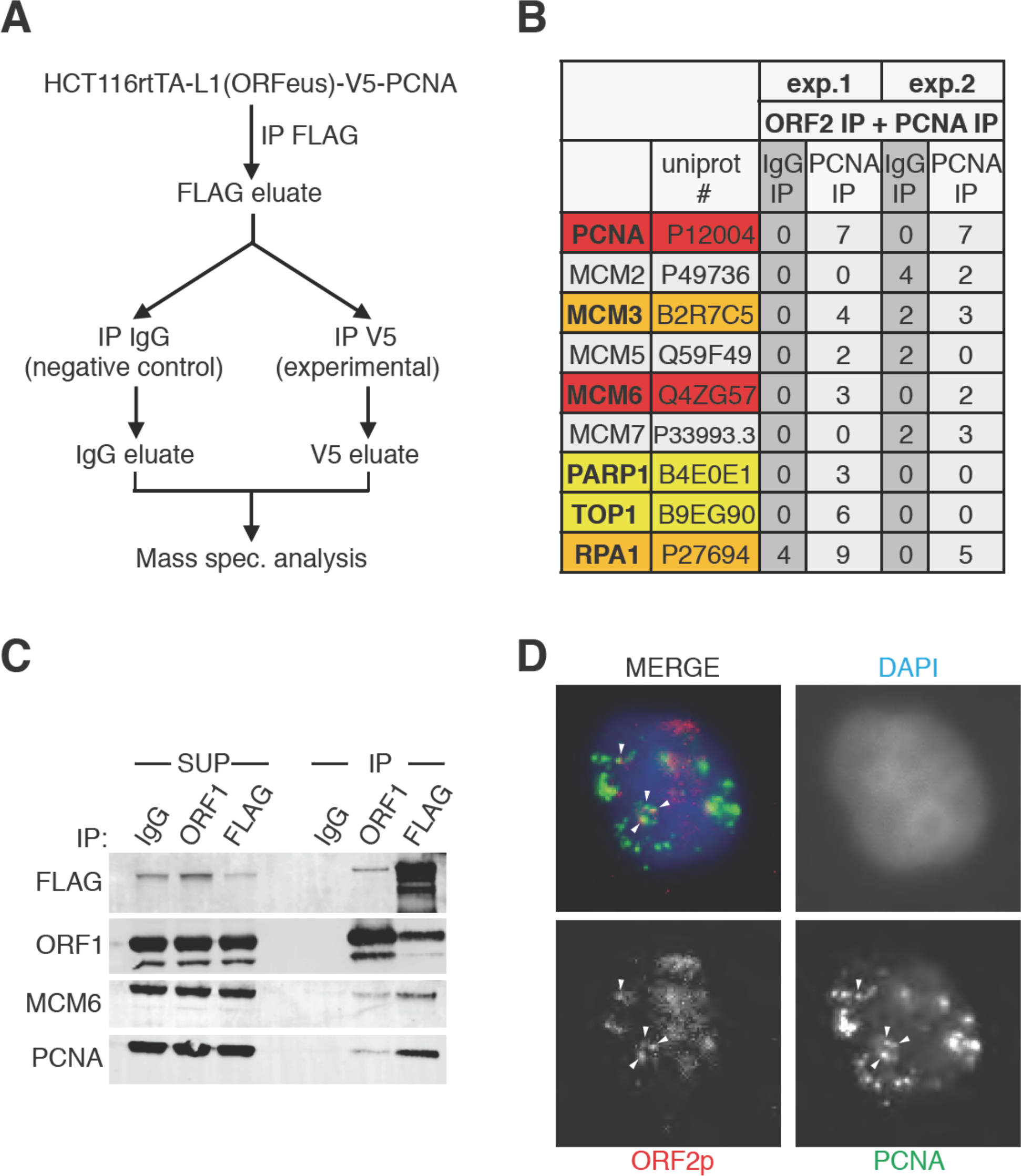
ORF2p is on replication forks with PCNA during S phase. (A) Schematic of sequential ORF2-PCNA immunoprecipitation and mass spectrometry analysis. HCT116 expressing V5-PCNA, rtTA and a recoded L1 (ORFeus) with flag-tagged ORF2p were used to immune-precipitate ORF2p. The immunoprecipitated complexes eluted with FLAG peptides were split in two and used for immunoprecipitation with V5 or IgG control antibodies. After native elution with V5 peptides the samples were analyzed by mass spectrometry. (B) Peptides numbers of known DNA replication fork proteins (identified by their UniProt number) obtained after mass spectrometry analysis of the ORF2p-PCNA/IgG sequential IP. Peptide counts of two independent experiments are reported. Red= high confident interactors that are found in both experiments and with no peptides in the IgG control IPs; orange= possible interactors that are found in both experiments and with some peptides in the IgG control IPs; yellow= low confidence interactors that are found in just one experiment and with no peptides in the IgG control IPs; gray= MCM proteins not identified as ORF2p interactors in our analysis. (C) Western blot of MCM6 and PCNA proteins co-immunoprecipitated upon IgG (control), ORF1 or FLAG(ORF2) IP. Grindates of 293T_LD_ cells expressing a recoded L1 with flag-tagged ORF2p were used. (SUP= supernatant after IP, IP=immunoprecipitation). (D) Immunostaining of ORF2p (red) and PCNA (green) in cells synchronized with double thymidine block and released for 2 h in S phase.

Collectively, our biochemical and proteomic work (Fig. 7) support the hypothesis that ORF2p is co-recruited with PCNA on sites of DNA replication during S-phase, and this fraction is most likely engaging in retrotransposition as demonstrated by our functional assays (Fig. 5).

## DISCUSSION

Despite the increasingly appreciated relevance of L1 retrotransposon to normal cellular physiology and disease etiology, many of the steps of L1 retrotransposon lifecycle in human cells are largely unknown. This lack of insight about L1 retrotransposons in human cells is unsurprising considering that many technical challenges hinder studies of this highly repetitive but poorly expressed element, which is effectively repressed by host somatic cells (Goodier, 2016) and therefore overexpression approaches can reveal otherwise hidden pathways.

It makes sense that nuclear localization of a L1 RNP particle comprising at least ORF2p, with EN and RT activity, bound to L1 mRNA, is essential for L1 retrotransposition to gain access to its target, genomic DNA. Despite this obvious observation, the nature of nuclear L1 RNPs and the process by which L1 gains entry into the nucleus are unknown. No functional nuclear localization signal (NLS) has been identified in the two L1 proteins ORF1p and ORF2p suggesting that their import into the nucleus is either mediated by interacting partners or by cellular processes such as the cell cycle and progression through mitosis during which the nuclear membrane breaks down, allowing the possible entrance of L1 RNPs into the nucleus. The former hypothesis is supported by several studies that show an essential role of cell division on retrotransposition and retrotransposition rate in tissue culture cells (Xie et al., 2013, Shi et al., 2007). On the other hand, other work showed the possibility of L1 retrotransposition in differentiated and non-dividing cells such as human neurons and glioma cells, albeit at substantially lower rates (Macia et al., 2016, Kubo et al., 2006). These seeming incongruities may be explained with a possible major mechanism of entry into the nucleus during mitosis and a less frequent mode of nuclear localization for L1 RNPs that is independent of the cell cycle and specific for some cellular state or cell type.

Through imaging, genetic and biochemical approaches, we show that L1 nuclear import as well as L1 retrotransposition has a strong cell cycle bias (Fig. 8). We also show that ORF1p, probably in a complex with ORF2p and L1 mRNA, enters the nucleus during mitosis, accumulating in cells in the G1 phase (Fig. 2). We observe a CRM1 mediated/leptomycin sensitive nuclear export of ORF1p (Fig. 3) that keeps the nuclear level of ORF1p low and helps explain the observation that ORF1p is always mostly cytoplasmic (Fig. 1 and Fig. 3). Future studies will need to explore the CRM1/ORF1p interaction and the role of this interaction in L1 RNP cellular dynamics and retrotransposition. The decoupling of the CRM1 role on the cell cycle from its importance on L1 cellular localization will be challenging but essential for the understanding of ORF1p nuclear export. We also show that even if L1 enters the nucleus in M phase, retrotransposition does not happen during cell division (M phase) but it is during the following S phase in which retrotransposition peaks (Fig. 4 and 5). The finding that L1 retrotransposition has a strong bias for S phase is, in retrospect, not entirely surprising considering that deoxynucleoside triphosphates (dNTPs), critically necessary for reverse transcription, are at high levels during the S phase and are greatly restricted during the other cell cycle stages (Hofer et al., 2012, Stillman, 2013). This layer of metabolic regulation may reflect an ancient adaptation to limit the proliferation of retroelements. dNTP concentration is tightly controlled by ribonucleotide reductase (RNR), the enzyme that converts ribonucleotide diphosphates (rNDPs) into dNDPs and by SAMHD1 (sterile alpha motif and HD-domain containing protein 1) that cleaves dNTPs to deoxynucleoside (Stillman, 2013). Indeed, SAMHD1 expression was found to restrict replication of lentiviruses such as HIV, by restricting availability of dNTPs (Hrecka et al., 2011, Goldstone et al., 2011). It is therefore not surprising that SAMHD1 was also shown to restrict LINE-1 retrotransposition (Zhao et al., 2013) and directly supporting the idea that dNTP concentration can profoundly limit L1 jumping. In complete accord with these observations is our finding that retrotransposition happens in S phase, during which dNTP concentration peaks, allowing efficient reverse transcription and thus L1 retrotransposition. This may thus be viewed as an adaptation of the retroelement to a host defense.

**Fig. 8.**
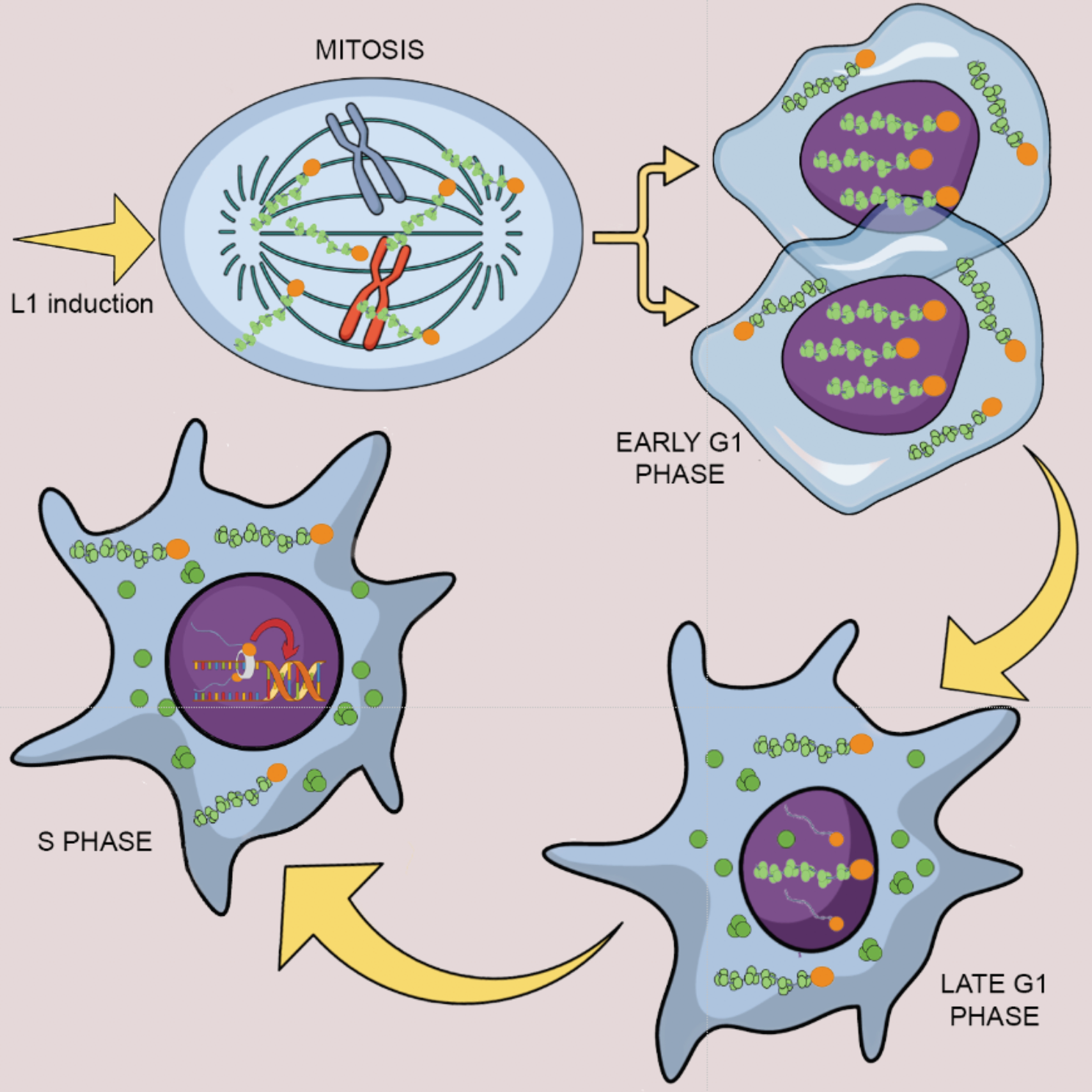
Model of L1 protein dynamics and L1 retrotransposition during the cell cycle. After induction of L1 expression, L1 RNPs formed by ORF1p (green balls), ORF2p (orange balls) and L1mRNA (blue line interacting with ORF2p), enter the nucleus as a result of mitotic nuclear membrane breakdown. In early G1 phase, when the nuclear membrane re-assembles, L1 RNPs are found in the nucleus. ORF1p, but not ORF2p is then exported from the nucleus through a CRM1 dependent mechanism leaving ORF2p and L1 mRNA into the nucleus. During S phase and DNA replication the ORF2p-mRNA L1 particle retrotransposes into new loci of the genome interacting with PCNA on the replication fork.

Our functional studies showing S phase bias of L1 retrotransposition are also corroborated by our biochemical observations that show that ORF2p is recruited to chromatin during S phase (Fig. 6) and suggest that ORF2p is recruited to a subset of sites of DNA replication with PCNA and MCM proteins (Fig. 7). Mass spectrometry analysis revealed interaction of the previously described PCNA/ORF2p complex (Taylor et al., 2013) with TOP1, RPA1 and PARP1 proteins (K.R. Molloy, 2017) all of which associate with replication forks. Interestingly, the PARP1 interaction suggest that L1 specifically interacts with stalled replication forks (Berti et al., 2013). Our co-localization of PCNA and ORF2p also supports the presence of ORF2p at potential sites of DNA replication, marked by PCNA staining during the S phase. PCNA and ORF2p immunofluorescence revealed that only some PCNA foci of replication overlap with ORF2p nuclear foci, suggesting that, at least in some instances, retrotransposing L1 may specifically interact with a subset of perhaps stalled replication forks. Conversely, not all ORF2p nuclear foci overlap with PCNA sites, suggesting that L1 interaction with the replication fork may represent just one of several modes used by L1 to select a DNA target site and retrotranspose.

Interestingly, our observations on ORF1p cytoplasmic/nuclear dynamic suggest deeper implications. The fact that ORF1p is exported from the nucleus before S phase, leads to the conclusion that retrotransposition, happening mainly during DNA replication, is mediated by RNPs depleted of ORF1p and constituted only or predominantly by ORF2p and L1 mRNA, a conclusion also supported by data presented by Molloy et al. (K.R. Molloy, 2017). In vitro studies of TPRT show that the first and presumably critical steps in retrotransposition can efficiently occur in vitro in the absence of ORF1p (Cost et al., 2002). The hypothesis that ORF1p is dispensable for the actual DNA cutting and reverse transcription steps in vivo, is supported by our observation that nuclear ORF1p can be specifically recognized by one of our antibodies (JH73g). We interpret this observation to mean that this antibody recognizes a specific conformational state of ORF1p unique to the nucleus. Not surprisingly, the nuclear form of ORF1p that is recognized by the JH73g Ab, has impaired binding to ORF2p, suggesting that once inside the nucleus, ORF1p may dissociate from the L1 RNPs destined to carry out the critical endonuclease/reverse transcription steps of retrotransposition during S phase. Moreover, most of the ORF1p does not interact with PCNA and MCM6 that, instead, interact mainly with ORF2p. We also observed (rare) instances of cells clearly expressing ORF2p in the nucleus in the absence of detectable ORF1p (Fig. 1 and (K.R. Molloy, 2017). Moreover, we observed clearly differential distribution of ORF2p, which forms nuclear puncta, and ORF1p, which has more diffuse localization, in the nucleus. This difference, though, could be due to the substantial differences in the amount of expressed protein between ORF1p and ORF2p. Finally, observations of HeLa.S-FUCCI cells expressing Halo tagged ORF1p show that ORF1p is never nuclear in cells in the S/G2/M phase. Halo tagged ORF2p, in contrast, was observed in the nucleus of certain cells in S/G2/M (fig. S7C) suggesting that, during these cell cycle stages, ORF2p is in the nucleus absent ORF1p. Overall these data suggest that chromatin bound and retrotransposing L1 particles are depleted of ORF1p and mainly consist of ORF2p in complex with L1 mRNA and host factors involved in retrotransposition. Future studies are necessary to better understand the differences of nuclear and cytoplasmic ORF1p and the molecular processes that may mediate ORF1p depletion from L1 RNPs. An attractive possibility is that ORF1p dissociation from L1 RNPs might be associated with delivering an ORF2-RNA RNP to chromatin, although we do not have direct evidence for this. It is tempting to imagine that in the nucleus, the absence of ORF1p trimers, thought to bind L1 mRNA every 50 nucleotides in the cytoplasm (Khazina et al., 2011), may promote ORF2p’s unhindered movement during reverse transcription of L1 mRNA in the process of TPRT. This hypothesis will need to be better explored in future studies, by examining ORF2p and L1 mRNA dynamics during the cell cycle. The lack of a sensitive Ab against ORF2p, the fact that most cells expressing ORF1p do not express ORF2p due to an unknown post-transcriptional mechanism controlling ORF2p expression (Taylor et al., 2013, Alisch et al., 2006, Luke et al., 2013) and the difficulties of detecting ORF2p and L1 mRNA even in the context of overexpression (Doucet et al., 2016), continue to technically challenge the study of L1 cellular dynamics.

More recent advances in the study of L1 retrotransposon, such as the construction and characterization of ORFeus with its increased expression and function (An et al., 2011, Han and Boeke, 2004), the use of smaller and brighter fluorescent tags that allow the exploration of the temporal axis of retrotransposition and the implementation of sensitive biochemical approaches (Sakaue-Sawano et al., 2008, Subach et al., 2009, Grimm et al., 2015) enabled us to discover new and unexpected interactions between L1 and the “host” cell. It is not surprising that retrotransposons, evolved within the human genome for millions of years, have “learned” to leverage important cellular pathways, such as the cell cycle and DNA replication, for their own purpose of spreading and increasing their genomic content (Boissinot et al., 2000, Boissinot and Sookdeo, 2016). On the other hand, it is also increasingly clear how cells respond to L1 expansions during evolution, engaging in an ongoing genetic arms-race (Daugherty and Malik, 2012, Molaro and Malik, 2016). For example, as previously proposed, the nuclear membrane may have represented one of many a barrier that retrotransposons had to overcome to maintain effective retrotransposition frequency (Boeke, 2003, Koonin, 2006).

## MATERIALS AND METHODS

### Cell lines

HeLa M2 cells (a gift from Gerald Schumann, Paul-Ehrlich-Institute; (Hampf and Gossen, 2007) were cultured in DMEM media supplemented with 10% FBS (Gemini, prod. number 100-106) and 1 mM L-glutamine (ThermoFisher/Life Technologies, prod. number 25030-081) (complete medium). Cells were routinely split in fresh medium upon reaching 80-90% confluency. During routine culture of the cells the medium was changed every 2-3 days.

293T_LD_ cells adapted to suspension (Taylor et al., 2013) were used for transfection with PEI and collected to generate cell grindates as previously described in (Taylor et al., 2013). Cell grindates were used for IPs presented in Fig. 7C.

HCT116 colorectal carcinoma cells were cultured in McCoy’s 5A media (Life Technologies/Gibco, prod. number 16600-108) supplemented with 10% FBS and 1 mM L-glutamine. A subline of these cell line and expressing a V5-PCNA protein was generated using a CRISPR approach. A single Cas9-gRNA vector (PM192) was generated by Golden Gate reaction (Ran et al., 2013) using vector pX459V2.0 (a gift from Feng Zhang, Addgene, plasmid number 62988) and an annealed DNA duplex: CACC G GAAAAGACTTCAGTATATGC

A gBlock DNA purchased from IDT integrated DNA technologies was used as donor DNA.

PM-GB3:

ccgtgggctggacagcgtggtgacgtcgcaacgcggcgcagggtgagagcgcgcgcttgcggacgcggcggcattaaacggttgcaggcgtagcagagtggtcgttgtctttctagGTCTCAGCCGGTCGTCGCGACGTTCGCCCGCTCGCTCTGAGGCTCGTGAAGCCGAAACCAGCTAGACTTTCCTCCTTCCCGCCTGCCTGTAGCGGCGTTGTTGCCACTCCGCCACCATG GGT AAG CCT ATC CCTAACCCTCTCCTCGGTCTC

GATTCTACGGGAGAAGGGCAAGGGCAAGGGCAAGGGCCGGGCCGCGGCTAC GCGTATCGATCCTTCGAGGCGCGCCTGGTCCAGGGCTCCATCCTCAAGAAGG TGTTGGAGGCACTCAAGGACCTCATCAACGAGGCCTGCTGGGATATTAGCTC CAGCGGTGTAAACCTGCAGAGCATGGACTCGTCCCACGTCTCTTTGGTGCAG CTCACCCTGCGGTCTG

The donor DNA was transfected together with PM192 plasmid using Fugene-HD reagent (Promega, Madison, WI; prod. number E2311) in HCT116 plated in a 6 well plate. Cells were transfected with 200 ng of donor DNA, 1.5 µg PM192 and 6.8 µl Fugene-HD in 100 µl Opti-MEM (Thermo Fisher scientific, prod. number 31985088). 24 h after transfection cells were selected in 1ug/ml puromycin for additional 48 h. Single clones were picked after serial dilution of the cells in complete media without puromycin. Clones were screened in 96 well plates for expression of V5 by immunofluorescence staining. The positive clones were then validated by genomic DNA PCR using primers flanking the site of CRISPR cut. The amplified band was gel isolated and sequenced by Sanger sequencing. The primers used are the following:

JB17488-PCNACseqF

CTGCAGATGTACCCCTTGgt

JB17489-PCNACseqR

GACCAGATCTGACTTTGGACTT

The positive clones were then also validated by western blotting analysis using an antibody against PCNA and V5 tag.

The HCT116-V5-PCNA cell line was then used to generate stable cell lines expressing the rtTA transactivator. The pTet-ON advance vector (Clonetech, prod. number 631069) was transfected and cells were then selected for several weeks using media supplemented with 250µg/ml neomycin. Clones were screened using a construct encoding GFP under the control of a tetracycline activated promoter. Finally, HCT116-V5-PCNA-rtTA cell lines stably maintaining episomal pCEP-puro-plasmids expressing ORFeus (LD401) were generated and cultured under puromycin selection (1µg/ml) to prevent the loss of the L1 plasmids and neomycin (250µg/ml) to prevent the loss of rtTA transactivator.

120 15cm plates of HCT116-V5-PCNA-rtTA-LD401 cells were used to immunoprecipitate ORF2p-Flag using Flag-M2 antibodies (Sigma, prod. number

F1804). The immunoprecipitated protein complexes were then used as input to immunoprecipitate V5-PCNA using V5 antibodies (Invitrogen, prod. number 46-1157). Immunoprecipitation assays were conducted as described below.

HeLa.S-Fucci cells were purchased from the Riken BRC cell bank (RIKEN BioResource Center, Japan, prod. number RCB 2812). Stable HeLa.S-FUCCI cells expressing rtTA were generated transfecting a pTet-ON advance vector (Clontech, prod. number 631069) subcloned to carry a blasticidin resistance cassette instead of a neomycin resistance cassette. Cells were selected for several weeks in 15 µg/ml blasticidin and several clones screened for the expression of firefly under a control of a doxycycline promoter (gift from S.K. Logan laboratory). The selected stable HeLa.S-FUCCI-rtTA cell lines were cultured in complete DMEM media supplemented with 15ug/ml blasticidin.

### Plasmids and DNA constructs

LD401 (ORFeus with 3xFlag ORF2) and MT302 (L1rp with 3xFlag ORF2 and L1 5’UTR) plasmids were previously described and characterized in (Taylor et al., 2013). PM160 (ORfeus with L1rp 5’UTR) plasmid was obtained through Gibson reaction of a PCR L1 5’ UTR DNA fragment from MT302 used as template.

PM226 (L1rp with 3xFlag ORF2 and without L1 5’UTR) was generated by ligation of a synthetic 5’ end of L1rp without UTR to the BsiWI-PmlI cut MT302 vector.

EA79 (untagged ORFeus with GFP-AI cassette) was constructed subcloning ORFeus under a Tet inducible promoter into a pCEP-4 plasmid (ThermoFisher scientific, prod. number V04450) in which the hygromycin resistance cassette was substituted with a puromycin resistance cassette, as previously described in (Taylor et al., 2013) (pCEP-puro plasmid).

The fluorescent timer FT-AI cassette was built with Gibson assembly using two synthetic DNA fragments (Quinglan Biotech and Twist Bioscience) ligated into the BstZ17I 3’UTR of untagged ORFeus (PM260). Sequences of the fast fluorescent timer mCherry variant are the same of plasmid pTRE-Fast-FT (a gift from Vladislav Verkhusha, Addgene plasmid number 31913)(Subach et al., 2009).

Constructs expressing Halotag7-ORF1p (PM285) and Halotag7-ORF2p (PM283) were made by Gibson assembly of a PCR DNA fragment encoding the HaloTag7 (pLH1197-pcDNA3.1-PfV-Halo, a gift from Liam Holt) and pCEP-puro-ORFeus vector. The tag was inserted right before ORF1p and ORF2p stop codon downstream of a G4S linker.

All constructs were verified by Sanger sequencing (Genewiz).

### Immunoprecipitation, electrophoresis and western analysis

HeLa M2 cells were lysed in SKL Triton lysis buffer (50 mM Hepes pH7.5, 150 mM NaCl, 1 mM EDTA, 1 mM EGTA, 10% glycerol, 1% Triton X-100, 25 mM NaF, 10 µM ZnCl_2_) supplemented with protease and phosphatase inhibitors (Complete-EDTA free, Roche/Sigma prod. number 11873580001; 1mM PMSF and 1mM NaVO_4_). NuPage LDS sample buffer (ThermoFisher Scientific, prod. number NP0007) supplemented with 350 mM b-mercaptoethanol was added to the samples before gel electrophoresis performed using 4-12% Bis-Tris gels (ThermoFisher Scientific, prod. number WG1402BOX). Proteins were transferred on Immobilon-FL membrane (Millipore, prod. number IPFL00010), blocked for 1hour with blocking buffer (LiCOR prod. number 927-40000):TBS buffer (50 mM Tris Base, 154 mM NaCl) 1:1 and then incubated with primary antibodies solubilized in LiCOR blocking buffer:TBS-Tween (0.1% Tween in TBS buffer) 1:1. Secondary donkey anti-goat antibodies conjugated to IRDye680 (anti-rabbit) or IRDye800 (anti-mouse) dyes (LiCOR prod. number 926-32210 and 926-68071), were used for detection of the specific bands on an Odyssey CLx scanner (LiCOR). 293T_LD_ grindates used in Fig. 7c were lysed in extraction buffer (20 mM Hepes pH 7.4, 500 mM NaCl, 1% Triton X-100) supplemented with protease and phosphatase inhibitors.

Immunoprecipitations were performed using 5-10ul of dynabeads conjugated to primary antibodies (Dynabeads Antibody Coupling kit, Life Technologies, prod. number 14311D) incubated with lysates for at least 1 hour nutating at 4°C. After 5 washes in lysis buffer the immunocomplexes were eluted either in sample buffer, shaken with beads for 10 minutes at 70°C or using 3xFLAG (Sigma, prod. number F4799) or V5 (Sigma, prod. number V7754) peptides shaken with beads for 1 hour at 4°C. After elution, supernatants were collected and b-mercaptoethanol added to a final concentration of 350 mM.

Antibodies against ORF1p used in this study are:

4H1= Mouse monoclonal antibody targeting amino acids 35 to 44 of human ORF1p (Rodic et al., 2014, Taylor et al., 2013, Doucet-O’Hare et al., 2015) (Millipore, cat. number MABC1152).
JH74=Rabbit monoclonal antibody raised against the C-terminus of human ORF1p (Doucet-O’Hare et al., 2015) (a gift from Dr. Jeffry Han)
JH73g=Rabbit monoclonal antibody raised against the C-terminus of human ORF1p and immuno-purified by Genscript from hybridoma cells (a gift from Dr. Jeffry Han). Please note that this antibody is distinct from JH73 (Taylor et al., 2013) (see Fig. S6).
4632= Rabbit polyclonal antibody against human ORF1p (a gift from Dr. Thomas Fanning)

We also compared the previously described JH73 antibody (Taylor et al., 2013) with the JH73g antibody (Fig. S6). These two antibodies were derived from the same rabbit immunization using a purified globular ORF1p C-terminus (Dr. Jeffry Han, unpublished data). We confirmed that JH73g Ab is distinct from JH73 Ab displaying clear differences in the light chain migration and base peak chromatogram (Fig. S6).

### Mass spectrometry analysis

Samples were reduced and alkylated with DTT (1 hour at 57°C) and iodoacetamide (45 minutes at room temperature). The samples were then loaded on a NuPAGE 4-12% Bis-Tris gel (ThermoFisher Scientific, prod. number WG1402BOX) and ran for only 10 minutes at 200V. The gel was then stained with GelCode Blue Staining Reagent (ThermoFisher, prod. number 24590) and the protein bands were excised. The gel plugs were cut into 1mm^3^ pieces, washed and destained with 1:1 (v/v) (methanol:100mM ammonium bicarbonate). After at least 5 solvent exchanges the gel plug was dehydrated by aspirating 100ul of acetonitrile (ACN) and further dried in the SpeedVac. In-gel digestion was performed by adding 250 ng of trypsin (ThermoFisher, prod Number 90057) onto the dried gel plug followed by 300 µl of ammonium bicarbonate. Digestion was carried out overnight at room temperature with gentle shaking. The digestion was stopped by adding 300 µl of R2 50 µM Poros beads in 5% formic acid and 0.2% trifluoro acetic acid (TFA) and agitated for 2 hours at 4°C. Beads were loaded onto equilibrated C18 ziptips and additional aliquots of 0.1% TFA were added to the gel pieces and the wash solution also added to the ziptip. The Poros beads were washed with additional three aliquots of 0.5% acetic acid and peptides were eluted with 40% acetonitrile in 0.5% acetic acid followed by 80% acetonitrile in 0.5% acetic acid. The organic solvent was removed using a SpeedVac concentrator and the samples reconstituted in 0.5% acetic acid. An aliquot of each sample was loaded onto an Acclaim PepMap100 C18 75-µm x 15-cm column with 3 µm bead size coupled to an EASY-Spray 75 µm x 50 cm PepMap C18 analytical HPLC column with a 2 µm bead size using the auto sampler of an EASY-nLC 1000 HPLC (Thermo Fisher Scientific) and solvent A (2% acetonitrile, 0.5% acetic acid). The peptides were eluted into a Thermo Fisher Scientific Orbitrap Fusion Lumos Tribrid Mass Spectrometer increasing from 5% to 35% solvent B (80% acetonitrile, 0.5% acetic acid) over 60 m, followed by an increase from 35% to 45% solvent B over 10 m and 45-100% solvent B in 10 m. Full MS spectra were obtained with a resolution of 120,000 at 200m/z, an AGC target of 400,000, with a maximum ion time of 50 ms, and a scan range from 400 to 1500m/z. The MS/MS spectra were recorded in the ion trap, with an AGC target of 10,000, maximum ion time of 60 ms, one microscan, 2m/z isolation window, and Normalized Collision Energy (NCE) of 32. All acquired MS2 spectra were searched against a UniProt human database using Sequest within Proteome Discoverer (Thermo Fisher Scientific). The search parameters were as follows: precursor mass tolerance ±10 ppm, fragment mass tolerance ± 0.4 Da, digestion parameters trypsin allowing 2 missed cleavages, fixed modification of carbamidomethyl on cysteine, variable modification of oxidation on methionine, and variable modification of deamidation on glutamine and asparagine and a 1% peptide and protein FDR searched against a decoy database. The results were filtered to only include proteins identified by at least two unique peptides.

Proteins identified exclusively in the V5-PCNA IP (total of 762 proteins identified) and not in the parallel IP with the control IgG (total of 535 proteins identified) were considered components of the PCNA-ORF2 complex. These proteins (279 proteins) were queried in the STRING database (Szklarczyk et al., 2015) to identify known protein-protein interactions between these candidates.

Mass spectrometry analysis of JH73 and JH73g antibodies was performed as followed:

the antibodies were buffer exchanged to 100mM ammonium bicarbonate using 7K molecular weight cutoff Zeba Spin Desalting columns (Thermo Scientific) as per the manufacturer recommended protocol to remove glycerol. Following buffer exchange, samples were denatured with 8M Urea in Tris-HCl solution. Denatured samples were reduced with dithiothreitol (2µl of a 1 M solution) at 37 ^0^C for 1 hour and then alkylated with iodoacetic acid at room temperature in dark for 45 minutes (12µl of a 1M solution). Samples were diluted to final urea concentration of 2M to facilitate enzymatic digestion. Each sample was split into two aliquots and digested using trypsin and pepsin. For trypsin digestion, 100ng of sequencing grade-modified trypsin was added and digestion proceeded overnight on a shaker at RT. To inactivate the trypsin, samples were acidified using trifluoroacedic acid (TFA) to final concentration of 0.2%. For pepsin digest, samples were acidified to pH <2 using 1M HCl and digested using 100ng of pepsin (Promega) for 1 hour at 37 ^0^C. Pepsin was heat inactivated by incubating the samples at 95 ^0^C for 15 minutes. Peptide extraction was performed by addition of 5µl of R2 20µm Poros beads slurry (Life Technologies Corporation) to each sample. Samples were incubated with agitation at 4 °C for 3 hours. Peptide extraction was performed as described above. Digested peptides were loaded onto the column using the HPLC set up described above. The peptides were gradient eluted directly into a Q Exactive (Thermo Scientific) mass spectrometer using a 30 minute gradient from 5% to 30% solvent B (80% acetonitrile, 0.5% acetic acid), followed by 10 minutes from 30% to 40% solvent B and 10 minutes from 40% to 100% solvent B. The Q Exactive mass spectrometer acquired high resolution full MS spectra with a resolution of 70,000, an AGC target of 1e6, maximum ion time of 120 ms, and a scan range of 400 to 1500 m/z. Following each full MS twenty data-dependent high resolution HCD MS/MS spectra were acquired using a resolution of 17,500, AGC target of 5e4, maximum ion time of 120 ms, one microscan, 2 m/z isolation window, fixed first mass of 150 m/z, Normalized Collision Energy (NCE) of 27, and dynamic exclusion of 15 seconds.

### Immunofluorescence staining, live imaging and nuclear/cytoplasmic fractionation

For immunofluorescence staining cells were grown on coverslips or chamber-slides (Nunc, prod. number 154534) coated with 10 µg/ml fibronectin (ThermoFisher, prod. number 33016-015) in PBS for 4 hours-over night at 37°C. After plating cells were treated with 0.1 µg/ml doxycycline to induce expression of L1 (ORFeus or L1rp). After induction cells were prefixed adding formaldehyde (Fisher, prod. number F79-500) 11% directly to the culture media to a final concentration of 1%. After 10 minutes at room temperature the media/formaldehyde mixture was discarded and cells were fixed for 10 minutes at room temperature with formaline 4% in PBS (Life Technologies, prod. number 10010-049). For PCNA/ORF2p immunofluorescence presented in Fig. 7 cells were fixed for 20m in cold methanol. Cells were then washed twice in PBS supplemented with 10mM glycine and 3 times in PBS. Cells were then incubated for at least 1 hour at room temperature in LiCOR blocking buffer (LiCOR prod. number 927-40000). Upon blocking, cells were incubated over night at 4°C with primary antibodies diluted in LiCOR blocking buffer. The next day cells were washed 5 times in PBS with 0.1% Triton-X 100 and then incubated in secondary antibodies (Invitrogen, prod. number A11029, A11031, A11034, A11036, A32733) for 1 hours in the dark at room temperature. Cells were then washed 5 times in PBS with 0.1% Triton-X100 and 3 times in PBS and then coverslips or chamber slides mounted using VectorShield mounting media with DAPI (Vectorlab, prod. number H1200). Pictures were taken using an EVOSFL Auto cell imaging system (Invitrogen).

Pictures of HeLa-S.FUCCI live cells expressing Halotag7 ORF1p and ORF2p were obtained incubating the cells for 15 minutes with 100 nM JF646 dye (a gift from Timothee Lionnet)(Grimm et al., 2015) in complete FluoroBrite DMEM media (ThermoFisher, prod. number A1896701). After incubation with the dye, the cells were washed twice in PBS before observation under the microscope. Live cell imaging was performed using an EVOS-FL auto cell imaging system with on stage incubator. Live cell images and Z stack movies presented in the supplemental items (SI 2-3) were obtained using an Andor Yokogawa CSU-x spinning disk on a Nikon TI Eclipse microscope and were recorded with an scMOS (Prime95B, Photometrics) camera with a 100x objective (pixel size 0.11µM). Images were acquired using Nikon Elements software and analyzed using ImageJ/Fiji (Schindelin et al., 2012).

Cytoplasmic/nuclear fractionation was performed using a protein fractionation kit (Thermo prod. number 78840).

### Cell synchronization

HeLa M2 cells were synchronized in M phase by nocodazole treatment and mitotic shake off. Briefly, cells were treated for 12 h with nocodazole 60 ng/ml and mitotic cells collected in the supernatant after vigorous tapping of the plate. Cells were then washed three times in complete media and released into the cell cycle in fresh complete DMEM media.

HeLa cells were synchronized in G1/S boundary by double thymidine synchronization. 0.35x10^6^ cells were plated in 10 cm culture plates in complete media. After six hours media was exchanged with complete DMEM supplemented with 2mM thymidine freshly solubilized. After 18 h, cells were washed three times in PBS and released into the cell cycle with complete media. After 9 h, 2 mM thymidine media was added a second time to the cells for additional 15 h. Cells were then trypsinized, washed twice in PBS and released into the cell cycle in complete DMEM media.

### Propidium Iodide staining and cell cycle analysis

About 1x10^6^ cells were collected in a 1.5 ml tube and fixed at -20°C with 70% cold ethanol for at least 24 h. Cells were then pelleted by centrifugation at 420 rcf for 5 m and resuspended in HBSS:Phosphate citrate buffer 1:3 (phosphate citrate buffer: 24 parts 0.2M Na_2_HPO_4_ and 1 part of 0.1M citric acid) supplemented with 0.1% Triton X-100. Cells were incubated in this buffer for 30 m at room temperature, subsequently pelleted and resuspended in propidium idodide (PI) staining buffer (20µg/ml PI, 0.5 mM EDTA, 0.5% NP40, 0.2 mg/ml RNase A in PBS). Cells were incubated in PI staining buffer for 2 h at 37°C in the dark. After incubation PI fluorescence was analyzed on an Accuri C6 flow cytometer (BD bioscience) to determine the percentage of cells in each stage of the cell cycle.

### Retrotransposition assay

Retrotransposition assays shown in Fig. 4B were performed using HeLa-M2 cells stably expressing plasmid EA79 (pCEP-puro-ORFeus-GFP-AI). The experimental design is reported in Fig. 4A. Briefly, HeLa M2 cells transfected with EA79 plasmid were selected for at least 5 days in medium containing 1 µg/ml puromycin to generate stable cell lines that maintain episomal EA79. Expression of recoded L1 was induced with 1 µg/ml doxycycline. Measurements of GFP positive cells were done trypsinizing the cells and resuspending them in FACS buffer (HBSS buffer supplemented with 1% FBS, 1 mM EDTA and 100 U/ml of Penicillin-Streptomycin). To exclude dead cells from the measurement of GFP^+^ cells, 5 µg/ml propidium iodide (PI) was added for at least 5 minutes to the solution containing cells before measuring the percentage of GFP^+^ cells with an Accuri C6 flow cytometer (BD bioscience). GFP and PI signal were compensated before analysis. Retrotransposition assays presented in Fig. 4E-Fwere performed using HeLa M2 cells transfected with EA79 plasmid and always selected for 5 days in media containing 1 µg/ml puromycin as specified in the experimental design reported in Fig. 4C.

### Cell cycle analysis of retrotransposition using FT-AI reporter

Measurements of the cell cycle stage of cells that underwent retrotransposition using a fluorescent timer-AI reporter (PM260 plasmid) reported in Fig. 5B and C were done following two different approaches:

1. Reported in Fig. 5B and Fig. S9 A-B= HeLa M2 cells transfected with plasmid PM260 and selected for 5 days in puromycin were treated with doxycycline 1µ g/ml. After 24 hours of doxycycline induction, blue^+^red^-^ and blue^-^red^-^ (negative control) cells were sorted using a Sony SH800 sorter (Sony biotechnology Inc.). Cells were sorted directly into a tube containing cold 75% ethanol. The sorted cells were then stained with PI and their cell cycle stage determined as reported above.
2. 2) Reported in Fig. 5C and Fig. S9 C-D= HeLa M2 cells transfected with plasmid PM260 and selected for 5 days in puromycin were treated with doxycycline 1µ g/ml. After 24 hours of doxycycline induction, cells were incubated with complete media containing Syto61 DNA binding dye (ThermoFisher, prod. number S11343) and doxycycline 1µg/ml for one hour. Cells were then washed twice in PBS, trypsinized and resuspended in FACS buffer (see above). Fluorescence emissions from the fluorescent timer-AI reporter and Syto61 DNA dye were measured using an LSRII UV analyzer flow cytometer (BD Bioscience). The cell cycle stage of blue^-^red^-^ (negative control), blue^+^red^lower^ and blue^+^red^higher^ cells was determined.

Analysis of the percentage of cells in the different cell cycle stages was determined using FlowJo v10.2 software.

### Western blot quantification

Quantification of western blot protein bands was performed using Image Studio ver. 3.1 software on LiCOR CLx scanned images.

### Quantification of ORF1p and ORF2p cellular localization

Quantification of cellular localization of ORF1p and ORF2p was performed counting more than 1000 cells in three different cell preparations. For each experiment, LED intensity, camera gain and exposure were kept constant and set that background fluorescence was negligible using a negative control sample (cells not expressing L1) that will have no signal using the chosen setting.

For quantification reported in Table 3a, HeLa cells expressing ORFeus were plated on 8 wells chamber, induced with doxycycline, fixed and stained as described in the method section. Images of cells stained with JH74 antibody against ORF1p and Alexa 647 conjugated secondary antibody (Invitrogen, prod. number A32733), were collected with the Arrayscan VTI and quantified using the Compartmental Analysis Bioapplication. Briefly, 16 fields per treatment were acquired at 20x magnification and 2x2 binning (1104x1104 resolution). DAPI positive nuclei were identified using the dynamic isodata thresholding algorithm after minimal backgrounds subtraction. DAPI images were used to identify cell nuclei and to delineate the nuclear edges (x=0). A “circle” smaller than the nucleus (x=-4) was used to identify cells with nuclear ORF1p and a “ring” outside the nucleus (1<x>8) was used to identify cells expressing cytoplasmic ORF1p. Limits of fluorescence were set so that no cells were considered positive for preparations of cells not treated with doxycycline (negative control). The reported parameters are explained below:

Total= total number of DAPI nuclei counted;

Nuclear ORF1p= cells with fluorescence signal inside the circle (x=-4) higher than the limit (higher fluorescence signal of the circle in the negative control) Cytoplasmic/total ORF1p= cells with fluorescence signal inside the ring (1<x>8) higher than the limit (higher fluorescence signal of the ring in the negative control).

Because all cells expressing L1 show expression of ORF1p in the cytoplasm, the number of cells with cytoplasmic ORF1p was considered as the number of total cells expressing ORF1p. The percentage of cells with nuclear ORF1p was calculated as: (number of cells with nuclear ORF1p) *100/(number of cells with cytoplasmic ORF1p=total number of cells expressing ORF1p).

All results are reported as mean and the error calculated as standard deviation (S.D.) or standard error of the mean (S.E.M). Data are considered to be statistically significant when p < 0.05 by two-tailed Student’s T test. In Fig. 4, asterisks denote statistical significance as calculated by Student’s T test (^*^ p < 0.05; ^**^p < 0.01; ^***^p < 0.001).

### Proximity analysis

HeLa M2 cells expressing ORFeus for 24 hrs were plated on fibronectin treated chamber-slides, treated with 0.1 µg/ml doxycycline, fixed and stained using JH74 rabbit primary antibody against ORF1p, Alexa 647 conjugated anti-rabbit secondary antibody and DAPI as described in the method section. Images were collected with the Arrayscan VTI system and cell positions within each slide obtained using the Compartmental Analysis Bioapplication. To quantify if cells with nuclear ORF1 are significantly closer to each other compared to random cells, we first calculated the shortest distance for each nuclear ORF1 cell to another nuclear ORF1 cell. Next, we randomly and repeatedly (n=1000) select the same number of cells as there are nuclear ORF1 cells and obtain the distribution of distances that correspond to random localization of non-nuclear ORF1 cells. We used the random distribution to calculate the p-value and a false discovery rate (FDR). Finally, we compared the distribution of distances for nuclear ORF1 cells that are significantly closer to each other (p-value=0.1) with the distribution of a random sample that did not express nuclear ORF1 using a Wilcoxon rank sum test.

## DATA AND SOFTWARE AVAILABILITY

Mass spectrometry data for ORF2p and V5-PCNA sequential IP presented in Fig. 7 have been deposited in MassIVE archive under submission number MSV000081124 and in Proteome Xchange archive under submission number PXD006628.

## ACKNOWLEDGMENTS

This work was supported by NIH grant P50GM107632 to J.D.B. The cytometry and cell sorting, High Throughput Biology (HTB) and Proteomic cores are partially supported by Laura and Isaac Perlmutter Cancer Center Support Grant, (NIH/NCI P30CA16087) and NYSTEM Contract C026719 (HTB core) and NIH/ORIP 1S10OD010582 grant (proteomic core). We thank Gregory Brittingham and Dr. Liam Holt for their help in collecting confocal images and movies.

## SUPPLEMENTAL INFORMATION

**Figure S1.**
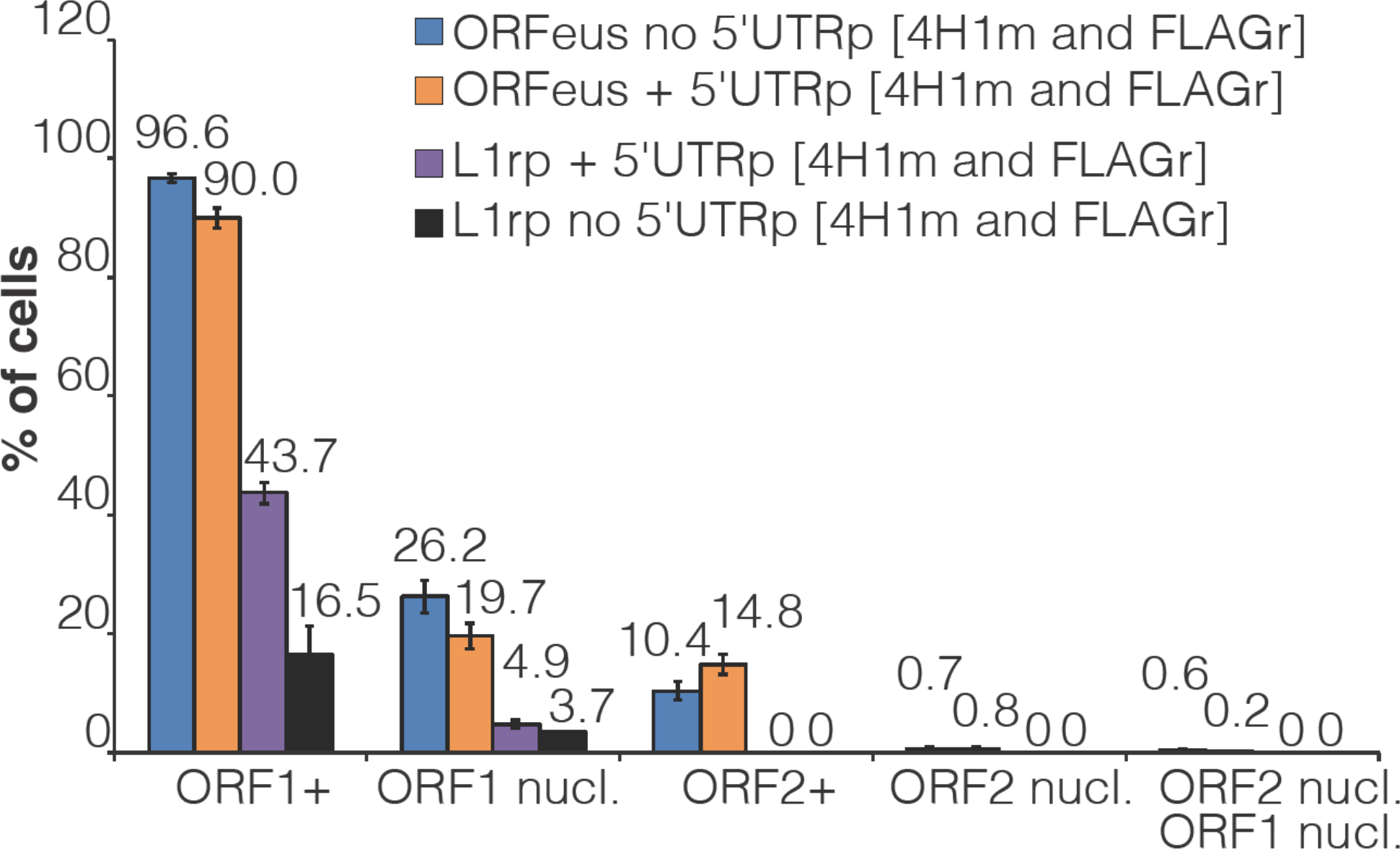
Role of L1 5’UTR in ORF1p and ORF2p localization. Quantification of nuclear and cytoplasmic ORF1p and ORF2p in HeLa M2 cells expressing the indicated recoded or non-recoded L1s with or without the 5’UTR. The percentage of cells is reported over each column bar. The error (standard error of the mean= s.e.m.) was calculated from at least 10 different fields (20X) containing at least 100 cells.

**Figure S2.**
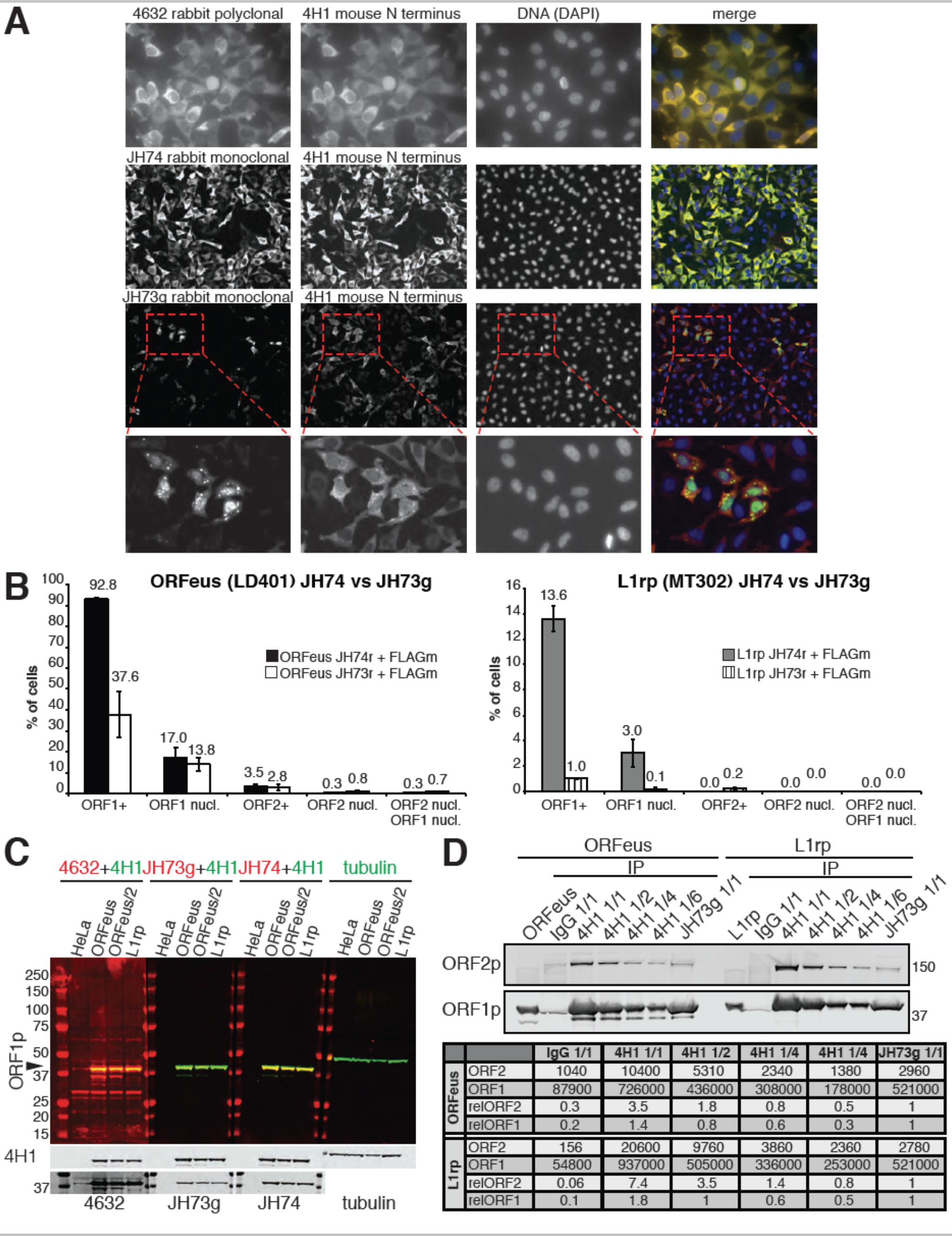
ORF1p antibody comparison. (A) Comparison of immunofluorescence stainings using several antibodies against ORF1p and HeLa M2 cells expressing a recoded L1. 4H1 mouse antibody (red) was compared against co-staining of rabbit 4632, rabbit JH74, rabbit JH73g antibodies (all shown in green). DAPI staining (blue) was used to label the nucleus. The lower row of pictures shows a magnification of clustered cells expressing nuclear ORF1p and stained with 4H1 and JH73g antibody. (B) Quantification of nuclear and cytoplasmic ORF1p and ORF2p in HeLa cells expressing a recoded L1 (ORFeus) or a native L1 (L1rp) and stained with JH74 or JH73g antibodies in combination with mouse anti-FLAG M2 antibody. The error is expressed as S.E.M. calculated from at least 10 different fields (20X) containing at least 100 cells. (C) Western blot of HeLa M2 cells expressing ORFeus, L1rp or no L1 (HeLa) blotted with the indicated antibodies in combination with 4H1 always in green. The same blotting is presented in colors to better show the overlap of red (from rabbit Abs) and green (from mouse Abs) signal, and in black and white to better show the single bands. Tubulin was used as loading control. 50 µg protein was loaded in HeLa, ORFeus and L1rp lanes and 25µg protein in the ORFeus/2 lanes. (D) Western blotting of ORF1p and ORF2p immunoprecipitated (IP) form lysates of HeLa cells expressing recoded (ORFeus/LD401) or native (L1rp/MT302) L1. Immunoprecipitation was conducted using IgG control, 4H1 or JH73g conjugated dynabeads. For IPs performed with 4H1 Ab several amounts of immunocomplexes were loaded on the gel: 1/1= same amount loaded for IgG, 4H1 and JH73g IPs; ½=half of the 1/1 amount; ¼=one fourth of the 1/1 amount; 1/6=one sixth of the 1/1 amount. Quantification of each band is reported as well as the relative quantification setting the intensity of JH73g IP band as 1.

**Figure S3.**
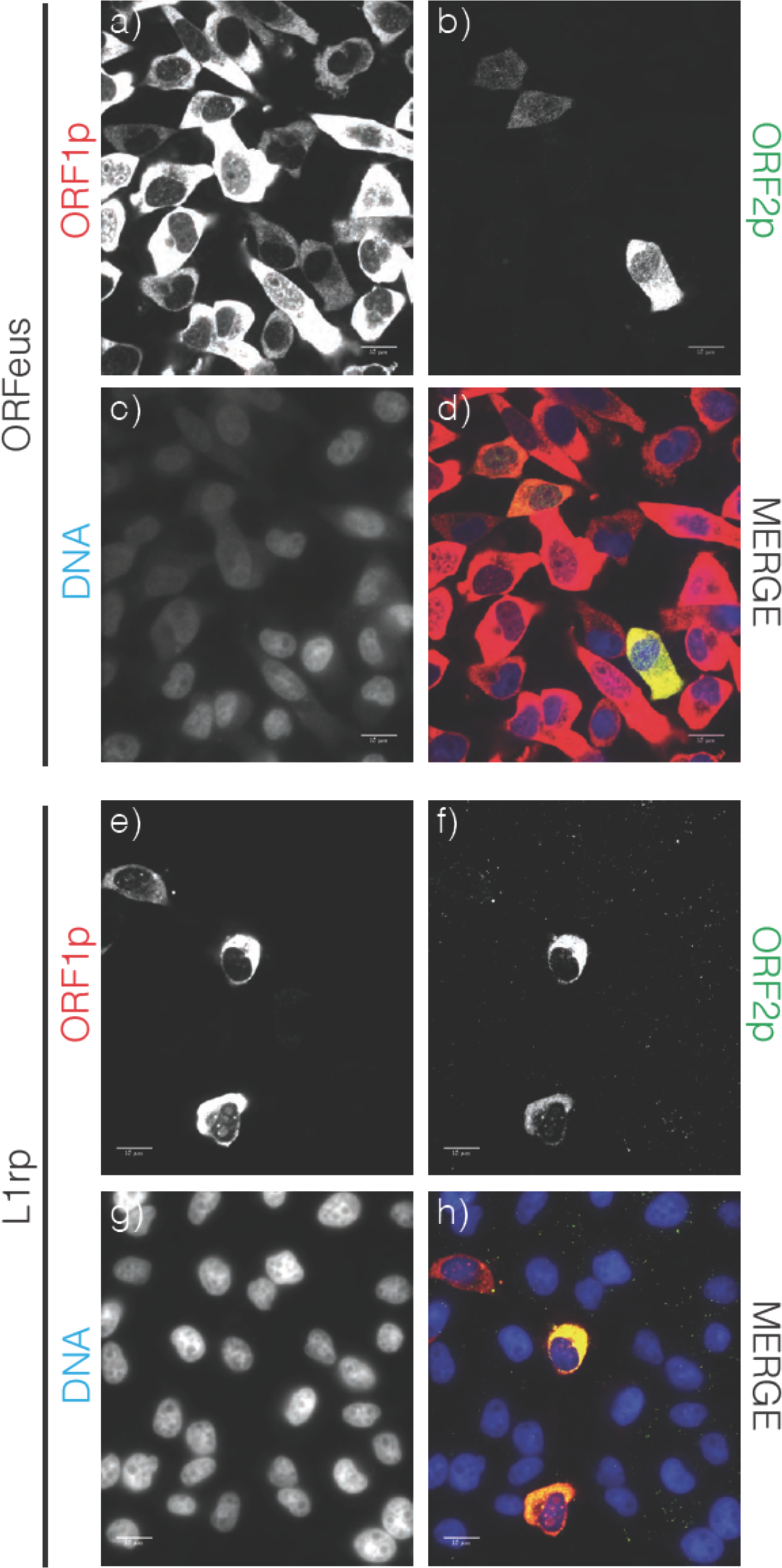
Confocal images of LINE-1 ORF1p and ORF2p. HeLa-M2 cells expressing recoded L1/ORFeus (A-D) or L1rp (E-H) cultured for 24 h in the presence of doxycycline 0.1 µg/ml. Cells expressing ORFeus are fixed with formalin and stained with 4H1 mouse anti-ORF1p (red) and rabbit anti-FLAG antibodies (green). Cells expressing L1rp are fixed and stained with JH74 rabbit anti-ORF1p (red) and mouse anti-FLAG-M2 (green). DNA is stained with Hoechst 33258. Complete Z-stack for the ORF1p and ORF2p images presented in this picture are shown in Supplemental item S4 (ORFeus) and S5 (L1rp).

**Figure S4 (movie). ORFeus ORF1p/ORF2p Z stack.**Z-stacks of recoded L1 (ORFeus) ORF1p (left) and ORF2p (right) expressed in HeLa-M2 treated for 24h with doxycycline 0.1 µg/ml. Red arrows points to cells expressing cytoplasmic and nuclear ORF2p and ORF1p; the green arrow points to a cell expressing cytoplasmic and nuclear ORF2p and just cytoplasmic ORF1p.

**Figure S5 (movie). L1rp ORF1p/ORF2p Z stack.**Z-stacks of L1rp ORF1p (left) and ORF2p (right) expressed in HeLa-M2 treated for 24h with doxycycline 0.1 µg/ml.

**Figure S6.**
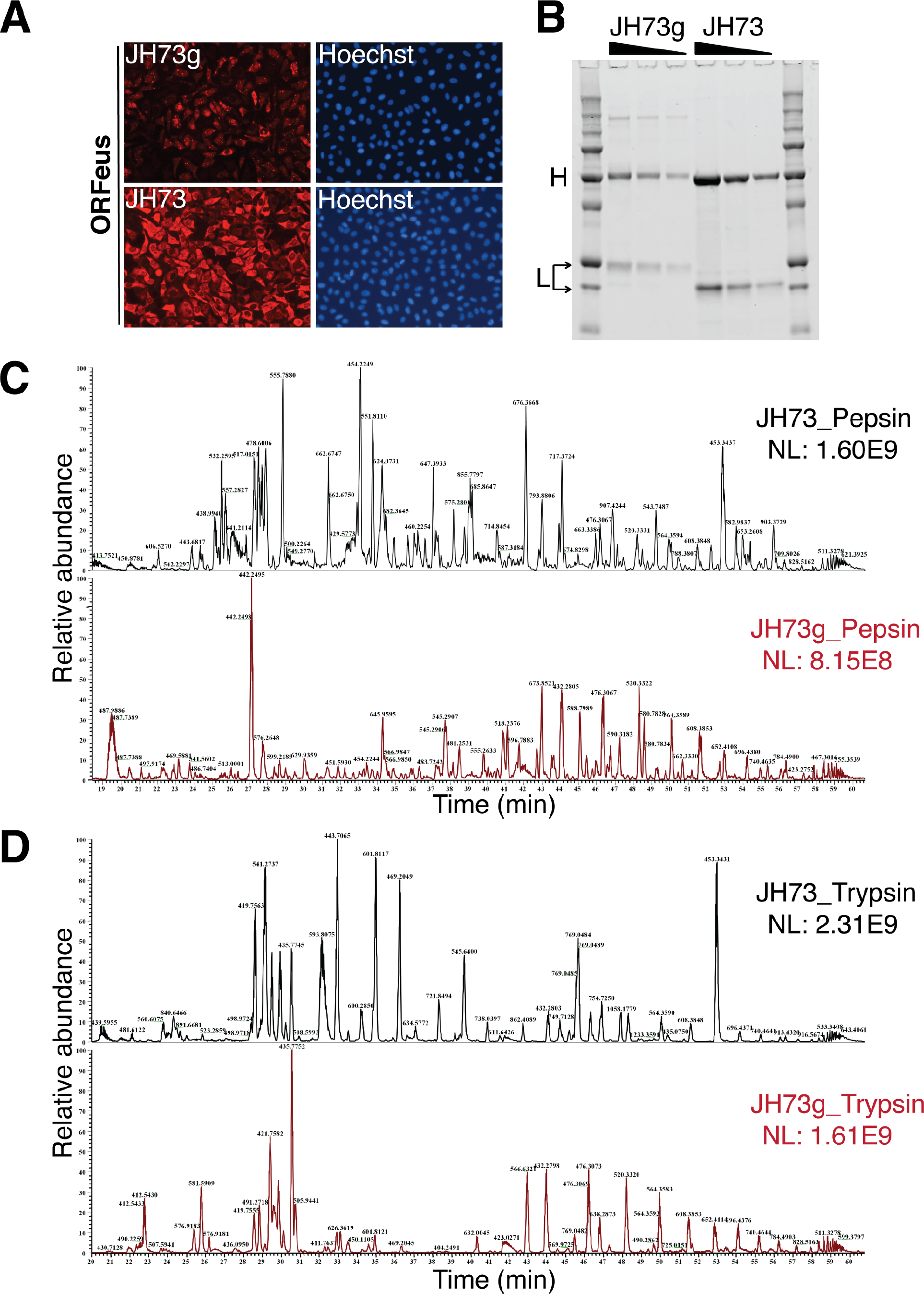
Comparison between JH73 and JH73g antibodies. (A) HeLa-M2 cells expressing recoded L1 (0.1 µg/ml doxycycline for 24h) stained with JH73g (top) or JH73 (bottom) and Hoechst 33258 (right panels). (B) SDS-PAGE of decreasing amount of JH73g and JH73 Abs. Overlay of base peak chromatogram of JH73 and JH73g antibodies digested using (C) pepsin and (D) trypsin. The base peak chromatogram plots the *m/z* value of the most abundant peptide ion eluting at a given time. NL represents the normalized ion intensity. For identical samples the major peptide ions in the chromatogram are similar. Here, there is little overlap between the major peptides in JH73 and JH73g, implying that the protein sequence and or the glycosylation pattern is different between the two antibodies.

**Figure S7.**
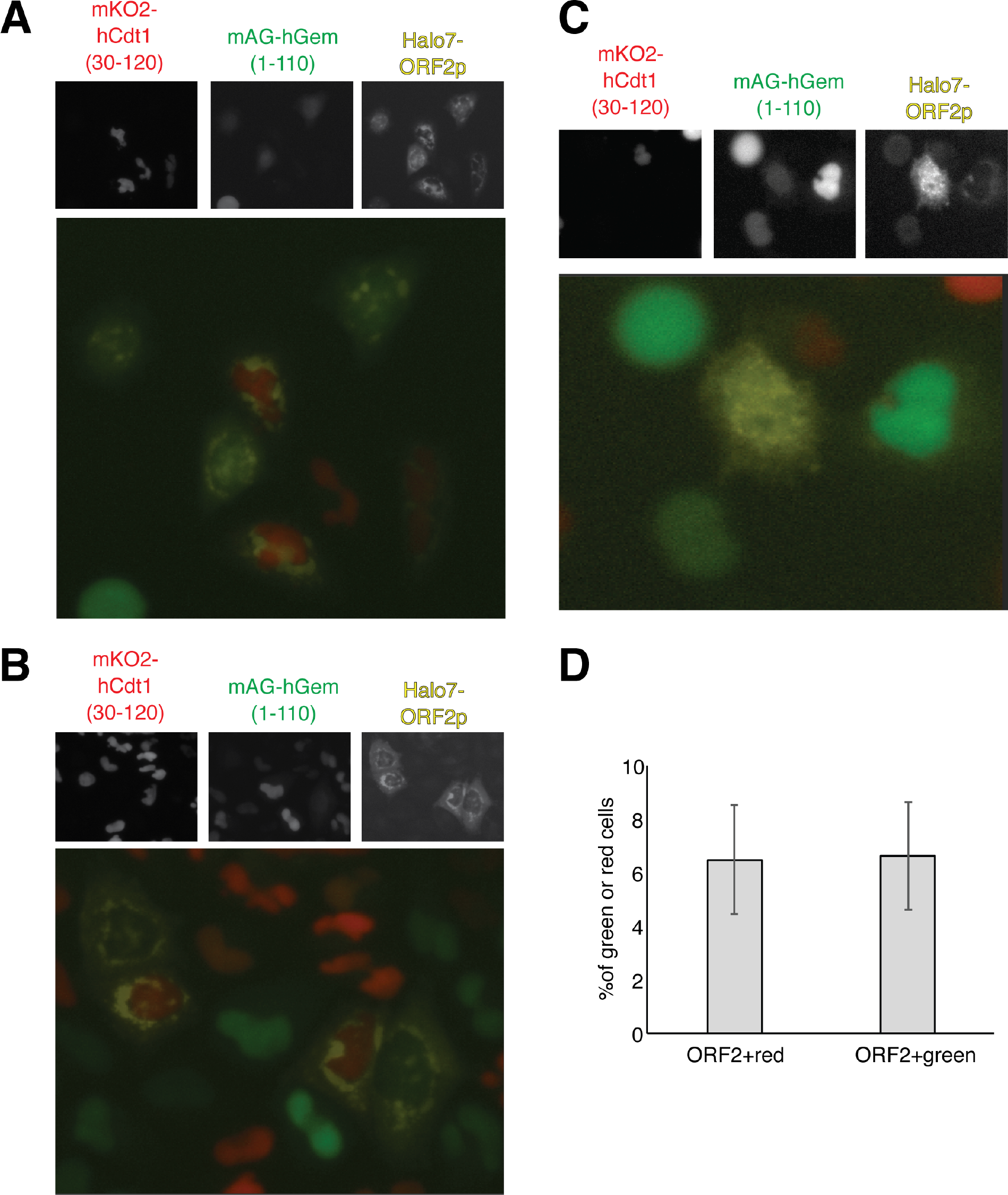
Expression and localization of Halo7-ORF2p in HeLa.S-FUCCI cells. AB) Representative pictures of HeLa.S-Fucci expressing rtTA and recoded L1 with an Halotag7 on the C-terminus of ORF2p. The human Cdt1 fragment (30-120) conjugated to mKO2 (monomeric Kusabira orange 2), human geminin fragment (1-110) conjugated to mAG (monomeric mutant of the green fluorescent protein Azami-Green) and the Halotag7-ORF2 were visualized using GFP (green), RFP (red) and cy5 (yellow) filters. Halotag7 was visualized using JH646 dye. The single emissions and the merged color picture are shown. An example of the cells with clear nuclear ORF2p is reported in (C). (D) quantification of cells expressing ORF2p (cytoplasmic or nuclear; ORF2^+^) and mKO2-hCdt1(30-120) (red) or mAG-hGem (1-110) (green) is reported. (error=S.D., n=3).

**Figure S8.**
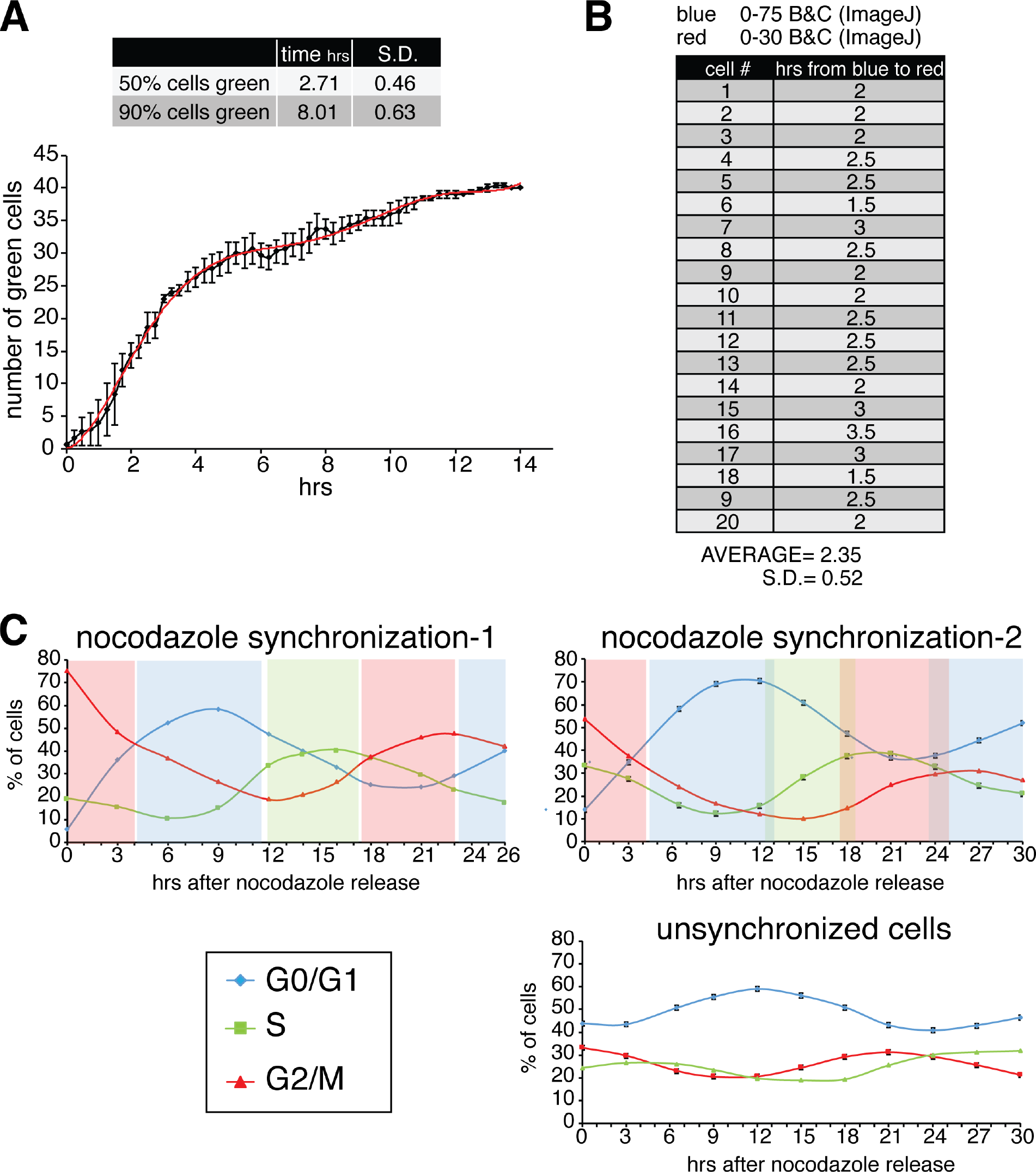
Dynamics of GFP expression, fast FT maturation and PI analysis. (A) Quantification of a fluorophore (GFP) expression under a Tet-CMV promoter. HeLa M2 cells expressing a pCEP-puro encoding GFP under doxycycline control, were used for a 15 h live cell imaging experiment. Time zero represents the time of doxycycline (1 µg/ml) addition to the medium. Pictures of the GFP signal were taken every 15 m using an EVOS-FL-auto with on-stage incubator. The R^2^ and equation of the best fitted curve are reported. (50% of the cells are green after 2.71±0.46 h; 90% of the cells are green after 8.01±0.63 h). (B) Quantification of fast-FT fluorophore maturation from blue to red. HeLa M2 cells expressing Tet-inducible fast fluorescent timer (FT) were used for 20 h live cell imaging measurements. Cells were pretreated with doxycycline for several hours before starting the collection of pictures every 30 minutes using DAPI (blue) and RFP (red) filters. 20 cells that started to express the FT protein within the timeframe of the experiment were analyzed (cell #). The brightness and contrast parameters (B&C) were set, as indicated, using ImageJ. The number of frames in which a cell showed blue signal but not red signal was counted and the time in which each cell transition from “only blue” to red was extrapolated. The average value and standard deviation (S.D.) for the 20 analyzed cells are reported in h (2.35±0.52 h). (C) HeLa M2 cells from Fig. 4 E-F were fixed and treated with propidium iodide (PI) for cell cycle analysis by cytofluorimetry. The % of cells in each cell cycle stage was measured for each time point. The shaded colored boxes indicate the cell cycle stage extrapolated from this analysis.

**Figure S9.**
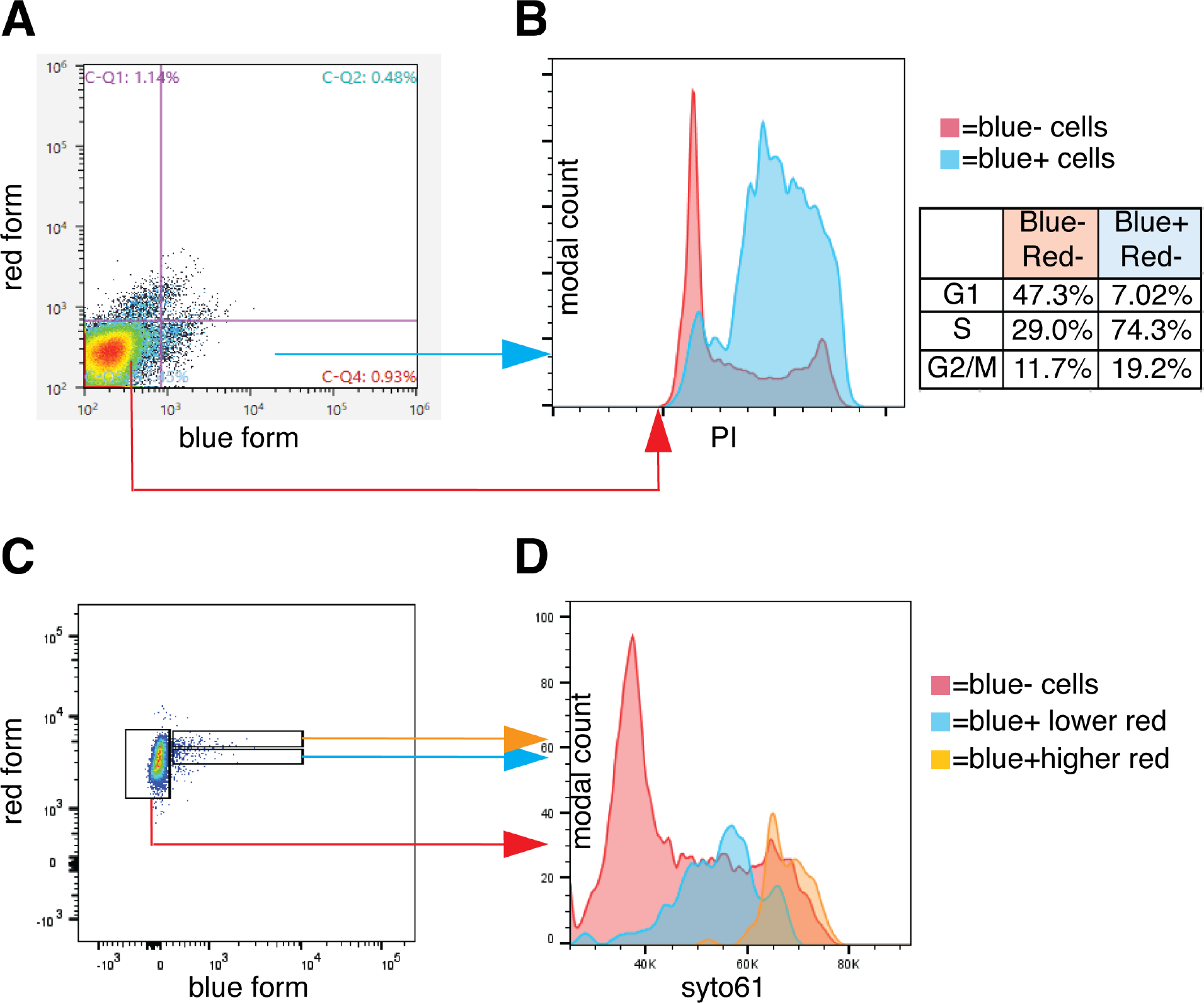
Cell cycle analysis of FT-AI. HeLa cells expressing the FT-AI retrotransposition reporter were treated with doxycycline 1ug/ml for 24 hrs. Upon treatment blue only cells (quadrant C-Q4) and double negative cells (quadrant C-Q3) were sorted as shown in the dot plot shown in (A). Sorted cells were then stained with PI and re-analyzed to measure the percentage of cells in each cell cycle stage (histograms and table in (B). The profiles of the two PI histograms from double negative (red) and only blue (blue) cells were overlaid using a modal Y axis. The inferred percentage of cells in each cell cycle stage is reported in the table. (C) HeLa cells expressing the FT-AI retrotransposition reporter were treated with doxycycline 1 µg/ml for 24 hrs. Upon doxycycline treatment cells were treated with 5 µM Syto61 DNA labeling dye for 1 h at 37°C. Cells were then analyzed with a LSRII-UV fluorimeter. Three populations were gated and considered for cell cycle analysis: blue negative (blue^-^) cells (red), blue positive (blue^+^) with higher red fluorescence (orange) and blue^+^ with lower red fluorescence (blue). The three cell cycle curves are overlaid in (D).

**Figure S10.**
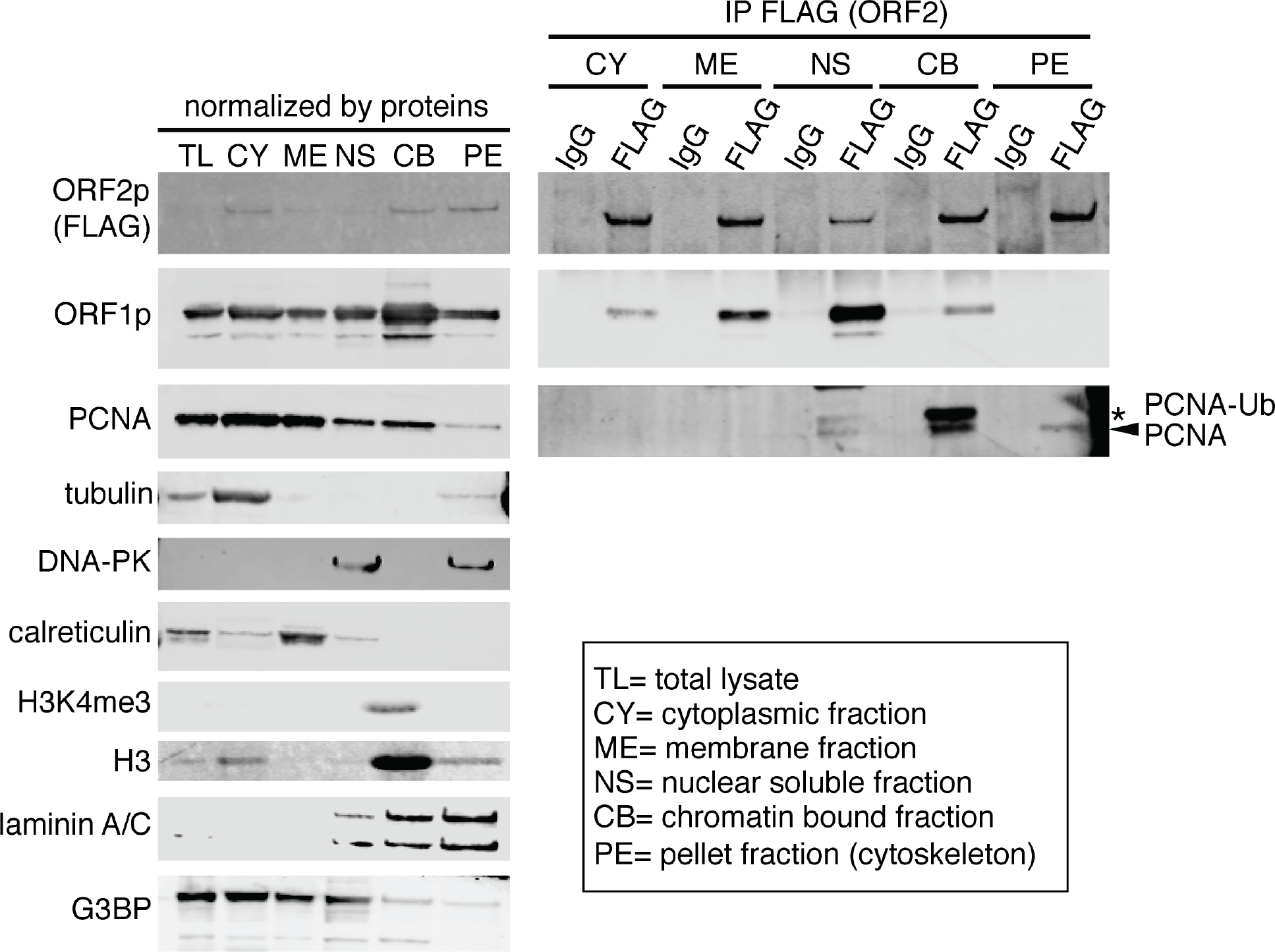
Analysis of nuclear fractions. HeLa M2 cells expressing a recoded L1 with a 3XFlagTag on ORF2p C terminus were processed for the isolation of protein cellular fractions using a protein fractionation kit (Thermo prod. number 78840). Upon fractionation ORF2p was immunoprecipitated (IP) from each fraction using FLAG-M2 (SIGMA) conjugated dynabeads. Western blotting analysis for each fraction was performed using the indicated antibodies. (tubulin= cytosol marker, DNA-PK= nuclear soluble marker, calreticulin= membranes marker, H3K4me and H3= chromatin marker, laminin A/C= nuclear membrane marker, GBP3=stress granules marker PCNAUb=ubiquitinated PCNA).

